# Spatiotemporal transcriptome at single-cell resolution reveals key radial glial cell population in axolotl telencephalon development and regeneration

**DOI:** 10.1101/2021.10.23.465550

**Authors:** Xiaoyu Wei, Sulei Fu, Hanbo Li, Yang Liu, Shuai Wang, Weimin Feng, Yunzhi Yang, Xiawei Liu, Yan-Yun Zeng, Mengnan Cheng, Yiwei Lai, Xiaojie Qiu, Liang Wu, Nannan Zhang, Yujia Jiang, Jiangshan Xu, Xiaoshan Su, Cheng Peng, Lei Han, Wilson Pak-Kin Lou, Chuanyu Liu, Yue Yuan, Kailong Ma, Tao Yang, Xiangyu Pan, Shang Gao, Ao Chen, Miguel A. Esteban, Huanming Yang, Jian Wang, Guangyi Fan, Longqi Liu, Liang Chen, Xun Xu, Ji-Feng Fei, Ying Gu

## Abstract

Brain regeneration requires a precise coordination of complex responses in a time- and region-specific manner. Identifying key cell types and molecules that direct brain regeneration would provide potential targets for the advance of regenerative medicine. However, progress in the field has been hampered largely due to limited regeneration capacity of the mammalian brain and understanding of the regeneration process at both cellular and molecular level. Here, using axolotl brain with extrodinary regeneration ability upon injury, and the SpaTial Enhanced REsolution Omics-sequencing (Stereo-seq), we reconstructed the first architecture of axolotl telencephalon with gene expression profiling at single-cell resolution, and fine cell dynamics maps throughout development and regeneration. Intriguingly, we discovered a marked heterogeneity of radial glial cell (RGC) types with distinct behaviors. Of note, one subtype of RGCs is activated since early regeneration stages and proliferates while other RGCs remain dormant. Such RGC subtype appears to be the major cell population involved in early wound healing response and gradually covers the injured area before presumably transformed into the lost neurons. Altogether, our work systematically decoded the complex cellular and molecular dynamics of axolotl telencephalon in development and regeneration, laying the foundation for studying the regulatory mechanism of brain regeneration in the future.

## INTRODUCTION

Brain is the most complex organ that controls emotion, memory, learning and many other functions, the brain in mammals, including human, has very limited regeneration capability, which even declines along development (Diotel et al., 2020; Kaslin et al., 2008; Tanaka and Ferretti, 2009), making researches or efforts on repairing of the injured brain extremely challenge. In contrast, some lower vertebrates such as teleost fish and salamanders preserve great ability of tissue regeneration (Joven and Simon, 2018; Kroehne et al., 2011; Lust and Tanaka, 2019; Maden et al., 2013). Among them, axolotl (*ambystoma mexicanum*), a tetrapod salamander species has been extensively studied (Amamoto et al., 2016; Echeverri and Tanaka, 2002; Gerber et al., 2018; Li et al., 2021a; Maden et al., 2013). Over a hundred years ago, the first forebrain regeneration in axolotls was observed in larvae (Burr, 1916). Similar regeneration phenotypes were documented as well in sex matured (adult) axolotls after removal of a large proportion of telencephalons (Amamoto et al., 2016; Maden et al., 2013). Amazingly, recent studies revealed that all lost cortical cell types, including neurons, could be reproduced at the lesion (Amamoto et al., 2016). As the telencephalon anatomy in axolotls and some other related salamander species is similar to that in mammals from the evolutionary point of view (González et al., 2017; Joven and Simon, 2018), in which the ventricular neural stem cells (NSCs) and neurons are located adjacent to the central lumen and peripheral pial surface respectively, Therefore, axolotls serve as an excellent models for studying the brain, in particular cortex regeneration, the discoveries of which may provide important insights in understanding the regeneration process in mammals.

Previous studies in varied regenerative species including axolotls have shown that ventricular radial glia cells (RGCs) respond to injury and contribute to brain regeneration (Berg et al., 2010; Joven and Simon, 2018; Kirkham et al., 2014). RGCs in adult salamanders are essentially the ancestor cells that give rise to nearly all cell types in the brain during early embryonic development (Merkle et al., 2004; Todd E. Anthony, 2004), and are maintained in the ventricular zone (VZ) (González et al., 2017). In contrast, most NSCs in mammals, except those in subventricular zone and hippocampal dentate gyrus, are almost completely consumed once the brain development is established. Under homeostatic state, dividing RGCs are sparsely located along the entire telencephalic VZ in axolotls, but only in a few confined VZ regions in red spotted newts. There are two groups of RGCs identified in telencephalon in red spotted newt, slow dividing and transient amplifying RGCs. The first group represents stem cell-like population, which express(Joven and Simon, 2018; Joven et al., 2018). Glial fibrillary acidic protein (GFAP) and glutamine synthetase, and show BrdU label-retaining property; the second group is located at the proliferative hot spots in VZ and frequently divide (Kirkham et al., 2014).

Upon injury, RGCs can be reactivated and expanded to broader areas in VZ. While both RGC groups could be detected close to lesion in red spotted newt, whether and how they contribute to brain regeneration is not clear (Kirkham et al., 2014; Maden et al., 2013). So far, only a few molecular signaling pathways that activate RGC and contribute to brain regeneration have been identified from salamanders and fish, such as Notch, FGF, and Gata3 (Kirkham et al., 2014; Kishimoto et al., 2012; Kizil et al., 2012). Interestingly, these signals are generally involved in brain development, implying that brain development and regeneration may share similarity in molecular regulation. Further investigation of this possibility and deeper understanding of brain regeneration requires more advanced technologies for systematic characterization of cell dynamics and the molecular expression in each cell type.

Many methods distinct in capturing strategy, resolution, and throughput, have been recently developed to obtain spatially resolved transcriptomic profiles of individual cells in a given tissue (Chen et al., 2021; Chen et al., 2015; Eng et al., 2019; Lubeck et al., 2014; Marx, 2021; Rodriques et al., 2019; Ståhl et al., 2016). However, one of the major technical challenges in the field is how to accurately assign a sequencing area to the physical location of each cell. Considering the complexity of the brain structure in general, well-defining of single cells will be essential to promote data accuracy and new discoveries. Mapping of the mouse brain or human cortex with spatial transcriptomics has been reported previously using barcoded slides (Maynard et al., 2021; Ortiz et al., 2020), but the resolution of these maps is limited by the diameters of the sequencing spots (100 μm or 55 μm, respectively). Data at such a resolution can only provide the average expression profiling for a mixture of over dozens of cells, which encumbers detailed investigation in regions with subtle networks of different cell types and molecular signals. In addition, the sequential fluorescence *in situ* hybridization (seqFISH+) (Eng et al., 2019) and multiplexed error-robust FISH (MERFISH) (Xia et al., 2019) have been developed for spatial profiling of gene expression of single cells, but their application may be restricted by the relative low throughput and the requirement of special equipment.

Taking advantages of a newly developed spatial-temporal transcriptomics approach—SpaTial Enhanced REsolution Omics-sequencing (Stereo-seq) (Chen et al., 2021) with the highest profiling resolution to date, we report here a *in situ* single-cell gene expression atlas and global cell dynamics in axolotl forebrain throughout development and the injury induced regeneration. We further identified the major brain RGCs participating in the pallium regeneration and establish molecular programs potentially involved in the activation of these cells and discovered presumably similar cell fate transition between telencephalon development and regeneration. Overall, our study provides comprehensive data sources for future investigation of the cellular and molecular mechanisms of brain regeneration.

## RESULTS

### Establishment of spatial transcriptome profile of axolotl brain at single-cell resolution

To first identify the individual cell types with precise location and transcriptome information in axolotl telencephalon, we prepared frozen sections of the adult axolotl telencephalon, followed by the Stereo-seq analysis on the entire section simultaneously (Chen et al., 2021) (Figure 1A). Considering the size of the axolotl cells (Herrick, 1948; Roth and Walkowiak, 2015; Westhoff and Roth, 2002), we collected coronal brain sections at 20 µm thickness to capture roughly a single-cell layer. As the Stereo-seq is based on DNA nanoball (DNB) sequencing technology (Porreca, 2010), for which each DNB spot on the chip is 220 nm in diameter and the center-to-center distance of two adjacent spots is 500 or 715 nm, we were able to capture transcripts at sub-cellular level (Figure 1A and Figure S1A).

**Figure 1.**
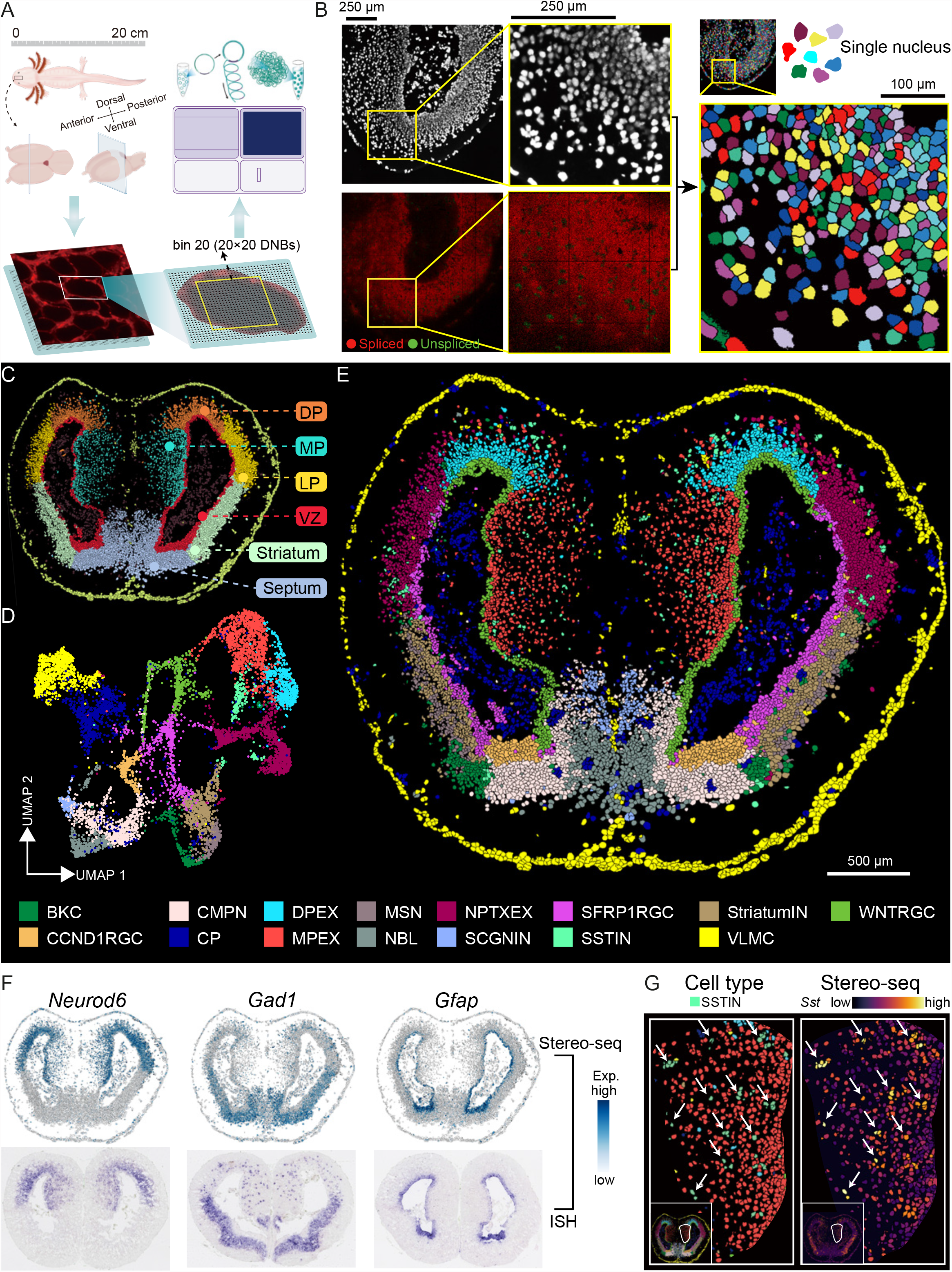
Stereo-seq visualizes spatial transcriptome profile of axolotl telencephalon at single-cell resolution. (A) Schematic diagram of Stereo-seq for axolotl telencephalon. *Step 1*, sample collection and frozen section preparation of the adult axolotl telencephalon. *Step 2*, *in situ* RNA capture from tissue placed on the chip. *Step 3*, cDNA amplification, library construction and sequencing. (B) Overlaying spatially assigned spliced- (red) and unspliced-transcripts (green) with corresponding ssDNA staining picture (left). Single-nucleus were defined based on ssDNA staining and represented by different colors (right). (C) Unsupervised spatially constraint clustering analysis considering location information of individually deduced nucleus at single-cell resolution. Cells were colored by region annotation. DP, dorsal pallium. MP, medial pallium. LP, lateral pallium. VZ, ventricular zone. (D) UMAP visualization of all cells profiled in adult axolotl telencephalon section. The colors correspond to the 15 identified cell types. (E) Spatial distribution of cell types identified in adult axolotl telencephalon section at single-cell resolution. Cell types are annotated by colored cubes. (F) Spatial visualization of selected gene expression on Stereo-seq map (top) and their correspondent *in situ* hybridization image (bottom). *Neurod6,* excitatory neuron marker; *Gad1,* inhibitory neuron marker; and *Gfap,* radial glial cell marker. (G) Distribution of SSTIN cells in the medial pallium region of adult axolotl telencephalon. Putative SSTIN cells are colored by green and signified by white arrow (left). Single cells highly express *Sst* gene are signified by white arrow (right). See also Figure S1-S4

We then attempted to define single cells on tissue sections by taking the advantage of the nucleic acid staining on cryosections to highlight the nucleus (Figure 1B), in which freshly transcribed mRNAs undergoing mRNA processing, including intron splicing are enriched (Gaidatzis et al., 2015; Gray et al., 2014; Zeisel et al., 2011). Indeed, intron-containing unspliced-transcript enriched areas overlapped nicely with stained nucleic acid signals, but are separated from spliced-transcript covered regions (Figure 1B, left panels), suggesting that the image of nucleic acid staining can be used to define the nucleus region. Inspired by this fact, we further employed watershed algorithm to our stereo-seq data to isolate the transcriptome in each DNA-staining defined area (Figure 1B, right panels), thus generating the spatial transcriptomic atlas of axolotl telencephalon at single-nucleus resolution. Each nucleus contains about 850 DNB spots, 6297 UMIs and 1682 genes on average (Table S1 and Figure S3 A-B). This high-resolution tool empowers us to delicately investigate diverse expression patterns of critical genes, with high similarity to *in situ* hybridization results (Figure S1B).

Using these data, we first performed unsupervised clustering analysis that considers both physical positions and global gene expression of individually deduced nucleus (details in methods). In total, we obtained six clusters of cells that show a patterned distribution on the brain section (Figure 1C), consistent with previous anatomical characterization of axolotl telencephalon, including ventricular zone (VZ), dorsal pallium (DP), medial pallium (MP), lateral pallium (LP), striatum and septum (González et al., 2017; Joven and Simon, 2018; Lazcano et al., 2021). To comprehensively dissect the cell type composition in the entire brain section, we next conducted unsupervised clustering analysis solely based on gene expression with Seurat (v4.0.2) (Hao et al., 2021). To this end, we categorized all identified nucleus into 15 cell types (Figure 1D) and further mapped them onto the telencephalon section according to the spatial information of each cell (Figure 1E). Cell identities were then determined by known marker genes in other species (Figure S2A; Table S2), and the spatial location of each cell type was further confirmed with the marker gene distribution results, which show great similarity to *in situ* hybridization (ISH) data (Figure S2B). For example, Stereo-seq based expression of classical excitatory neuron marker *Neurod6*, inhibitory neuron marker *Gad1* and RGC marker *Gfap* are almost identical to *in situ* hybridization results (Figure 1F).

As expected, all major brain cell types are present in distinct locations (Figure 1E and Figure S3C-E). Of note, there are three types of excitatory neurons enriched in pallium, named as dorsal pallium excitatory neuron (DPEX), medial pallium excitatory neuron (MPEX) and *Nptx*^+^ lateral pallium excitatory neuron (NPTXEX); In contrast, four types of inhibitory neurons, including striatum inhibitory neuron (StriatumIN), *Scgn*^+^ inhibitory neuron (SCGNIN), medium spiny neuron (MSN) and basket cell (BKC) are enriched in striatum or septum regions, physically separated from regions of excitatory neurons (Figure 1E). Remarkably, a fifth type of inhibitory neurons, *Sst*^+^ inhibitory neuron (SSTIN), were individually dispersed within the pallium regions, intermingled with excitatory neurons (Figure 1G and Figure S3E), consistent with previous study that *Sst* was expressed in scattered cells across DP and MP (Amamoto et al., 2016). Again, such exquisite distinction of cell types in space strongly endorsed the capability of Stereo-seq in realizing spatial transcriptome profiling of individual nuclei.

During brain development, neurons are formed by differentiation of neuron stem cells, or RGCs specifically for axolotl, which are also believed as the major contributing cell population for regeneration (Maden et al., 2013; Noctor et al., 2001). While it is known that axolotl RGCs mostly reside in the VZ region, we identified three clusters of cells located separately along the VZ regions with commonly high expression level of RGC specific markers genes, including *Sox2*, *Gfap*, *Nes*, *Vim*, *Fabp7* and *Slc1a3* (Figure S4), therefore named as *Wnt^+^* radial glial cell (WNTRGC) along the medial pallium side, *Sfrp1^+^* radial glial cell (SFRP1RGC) and *Ccnd1^+^* radial glial cell (CCND1RGC) according to their unique marker gene expression (Figure 1E). The distinct gene expression profile of each RGC type may suggest discrete functions. Indeed, CCND1RGCs highly express *Nes, Sox2,* cell cycle and ribosome related genes (Figure S2A; Table S2), suggesting they are potentially the progenitor cells for adult axolotl telencephalon maintenance (Barna, 2013; Bernal and Arranz, 2018; Calegari et al., 2005; Eming et al., 2014; Niu et al., 2015).

Other cell types identified included cholinergic, monoaminergic and peptidergic neuron (CMPN) and telecephalic neuroblast (NBL) in the septum, choroid plexus cells (CP) and vascular leptomeningeal cells in the out layer of the section (VLMC) (Figure 1E and Figure S2A). Altogether, with high-resolution stereo-seq, we provide a spatial cell atlas of axolotl telencephalon and transcriptome information for each cell, laying a cellular and molecular foundation for further development and regeneration studies. The interactive data portal can be browsed at https://db.cngb.org/stomics/artista.

### Cellular dynamics of RGCs throughout axolotl telencephalon development

It has been reported that in axolotls, RGCs in VZ region are responsible for brain development and regeneration, up on receiving stage-dependent developmental and injury cue (Amamoto et al., 2016; Maden et al., 2013). To more comprehensively understand cellular dynamics occurred during axolotl brain development, we carried out a series of spatial transcriptome analyses on sections of developmental (stage 44, 54, and 57), juvenile, adult, and metamorphosed axolotl forebrains (Figure 2A). Unsupervised clustering analysis based on gene expression with Seurat was applied to each section and in total 33 cell types were annotated coordinately across sections by their differentially expressed marker genes (Figure 2A, Figure S5A-C and S6; Table S1 and Table S2). In addition to the cell types identified in adult telencephalon (Figure 1), we also discovered 14 immature/intermediate cell types containing marker genes of both progenitor and differentiated cells (Figure 2A-B).

**Figure 2.**
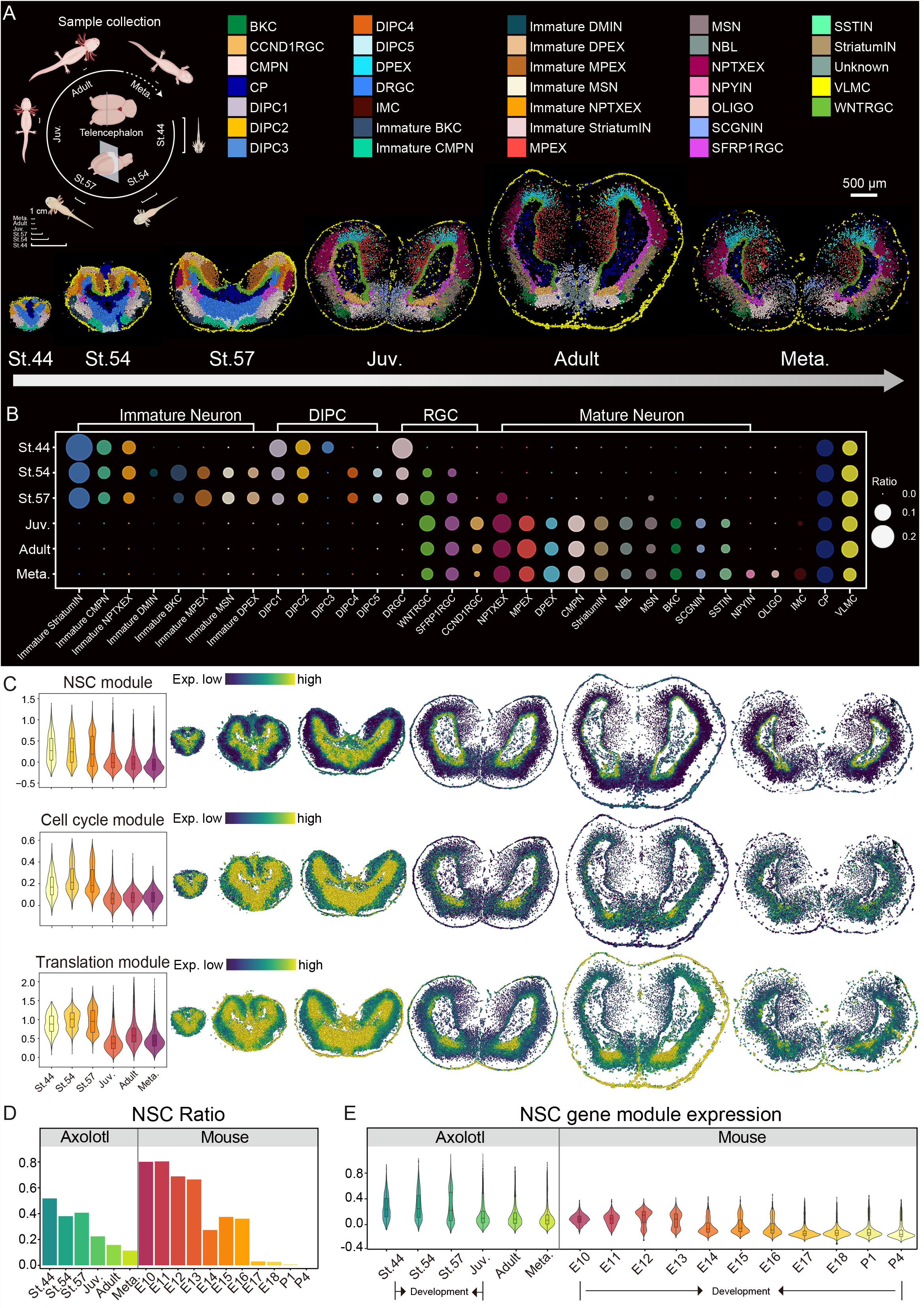
Spatio-temporal transcriptomic atlas of developmental, juvenile, adult and metamorphosis axolotl telencephalons at single-cell resolution. (A) Schematic diagram of sample collection (left). Unsupervised clustering of the axolotl telencephalon sections at stage 44 (St.44), stage 54 (St.54), stage 57 (St.57), juvenile (Juv.), Adult and metamorphosis (Meta.). Cell types are distinguished by colors. Scale bars, 500 μm (right). (B) Dotplot showing the ratio dynamic of cell types in the axolotl telencephalon from St.44, St.54, St.57, Juv., Adult and Meta. (C) Violin plot (left) and spatial visualization (right) of gene module expression defining NSC (neural stem cell), Cell cycle and Translation captured by Stereo-seq at different stages of axolotl telencephalon development. (D) Bar plot represents the cell ratio dynamic of NSC in the development process of axolotl and mouse. (E) Violin plot of gene module expression defining NSC at different stages in axolotl telencephalon development and in mouse embryonic development. See also Figure S5-S12

Most notably, we found a subpopulation of RGCs present in dominance throughout developmental stages, but were decreased in number and disappeared after juvenile stage (Figure 2A-B and Figure S7-S11). They expressed embryonic markers such as *Fzd5* and *Sox1*, and were named as development related RGCs (DRGCs). SFRP1RGCs, WNTRGCs and CCND1RGCs defined in adult telencephalon started to be detected since stage 54 and gradually became dominant RGC populations in designated locations from juvenile stage (Figure 2A-B). Along with DRGCs, immature neurons that expressed neuron lineage markers and *Stmn2*, *Tubb6, Dcx* were also detected at early developmental stages, the number of which progressively declined from stages 44 to 57 (Figure 2A-B and Figure S7-S11). Interestingly, the developmental intermediate progenitor cells (DIPCs) that co-expressed both RGC and immature neuron markers were discovered with similar temporal pattern as DRGCs and immature neurons (Figure 2A-B and Figure S7-S11), confirming the potential cell transition from RGCs to immature neurons as previously suggested (Noctor et al., 2001). In contrast to these developmental cell types, mature neurons, including subtypes of excitatory neurons and inhibitory neurons were gradually enriched, and the total number was increased from juvenile stage (Figure 2A-B and Figure S7-S11). This result nicely recapitulates the cellular dynamics of telencephalon development reported previously (Joven and Simon, 2018), suggesting that neurogenesis of axolotl telencephalon massively declines from juvenile stage, after which inactive SFRP1RGCs and WNTTGCs become WNTRGC made up the dominant population of RGCs, while DRGCs like CCND1RGCs only took up a small portion.

To further characterize the stemness and proliferation activity of RGCs in different VZ regions along the development, we analyzed the expression level of composite gene modules defining neural stemness, cell cycle and translation activity (methods, Table S3), all of which can help reveal the diverse aspects of stem cells (Figure 2C) (Fu et al., 2021; Temple, 2001). Overall, cells expressing high level of three gene modules were basically overlapped, distributed around the VZ, yet extended to the peripheral regions during early developmental stages, consistent with the fact of fast expansion of the brain size yet less neuron maturation in axolotl (Figure 2C) (Schreckenberg and Jacobson, 1975). Started from juvenile stage, cells positive for neural stemness, cell cycle and translation module gene expression significantly dropped in number and became restricted to the VZ region. Eventually, these active progenitor cells were enriched to the ventral area of the VZ region in adult, suggesting cells in this areas are responsible for adult brain maintenance similar to previous reports (Maden et al., 2013).

In contrast to the axolotl, the mouse has very limited regenerative ability upon brain cortex injury. To gain more insights into the molecular differences in stem cells between the axolotl and mouse brain, we compared the cellular and molecular dynamics of NSCs in mouse to that of RGCs in developing axolotl brains. Previously published single-cell RNA-seq data of developing mouse neocortex from Paola Arlotta lab were integrated with Stereo-seq data of developing axolotl telencephalons via Seurat (Figure S12 A-B) (Di Bella et al., 2021). Identified mouse NSCs and the combination of RGCs and DIPCs in axolotl were then compared. Interestingly, RGCs were relatively abundant in the VZ of axolotl brains, and their ratio sustained roughly even in adulthood (Figure 2D). Moreover, axolotl RGCs constantly expressed neural stemness module throughout the entire developmental and adult stages (Figure 2E). In contrast, NSCs in mouse brain gradually declined at later embryonic stages as well as after birth, accompanied with the degressive activity of neural stemness module (Figure 2D-E). These differences including the number and regeneration potential of stem cells between axolotl and mouse may partially explain the high regenerative capability of axolotl brains, which is absent in adult mammals.

### Injury specific RGCs contribute to pallium regeneration in axolotl

While axolotls harbor amazing regenerative capacity of the brain upon injury (Amamoto et al., 2016), the type of cells and their origins, as well as the underlying molecular events that direct the regeneration process and rebuild the exon network to recover the function are largely unknown. We next applied Stereo-seq to dissect the cellular and molecular dynamics during brain regeneration. Using a brain regeneration model established previously (Amamoto et al., 2016), in which a reproducible portion of the dorsal pallium in left telencephalic hemisphere of 11 cm length axolotl was removed by surgery, we collected brain tissues at 2, 5, 10, 15, 20, 30 and 60 days post injury (DPI) for Stereo-seq analysis, respectively (Figure 3A). Such efforts allowed us to investigate both immediate wound responses and chronic tissue regeneration process. By unsupervised clustering analysis based on gene expression with Seurat to each section (Figure S13A-D), we annotated cell types coordinately across sections by marker genes used in the developmental analysis (Figure S14). In total, 23 clusters were identified, including injury-specific cell populations that were not present during development (Figure 3A and Figure S13D). In line with previous reports that the injured site can be recovered to undetectable level in morphology in about 4 to 5 weeks (Amamoto et al., 2016), our data further revealed both cell types and their spatial distribution were basically recovered at 60 DPI in comparison with uninjured and the intact lateral side of injured brains (Figure 3A and Figure S13D).

**Figure 3.**
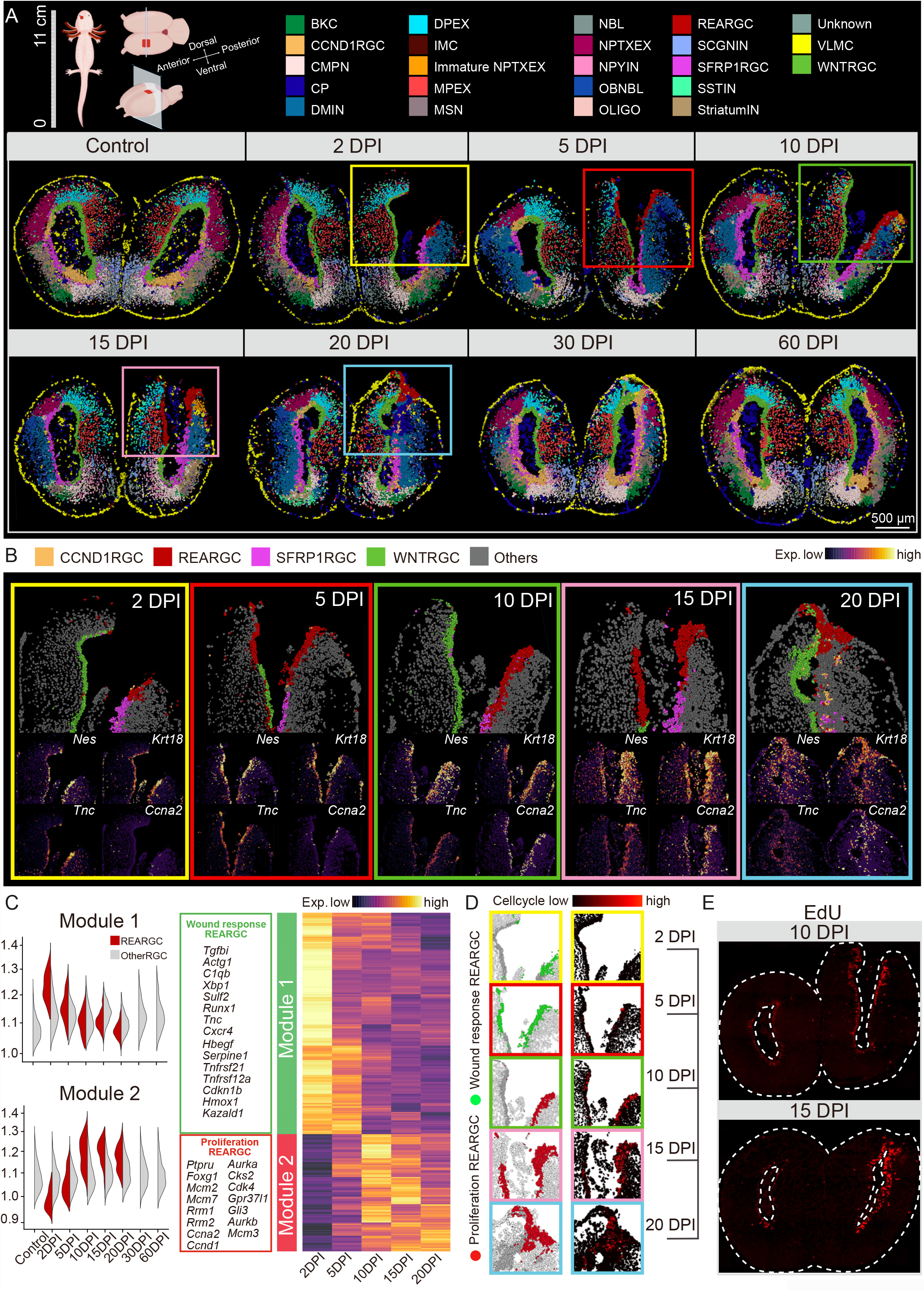
Spatiotemporal transcriptomic atlas of axolotl brain regeneration at single-cell resolution. (A) Schematic diagram of samples collection (upper left). Spatial visualization of cell types identified on the axolotl telencephalon sections at homeostatic and regenerative stages at single-cell resolution. Cell types are annotated by colored cubes on top. The squares indicate the regions analyzed in B and D. Scale bars, 200 μm. DPI, day post injury. (B) Spatial visualization of cell distribution in VZ zone (upper) and key marker expression (bottom) in the injured part of adult axolotl telencephalon section across 2DPI, 5DPI, 10DPI, 15DPI, 20DPI. Cell types are annotated by colored cubes on top. (C) Expression level of two gene modules in REARGC and other RGC cells from 2 DPI to 20 DPI (left). Heatmap reflecting the expression of genes from different modules (right). List of representative key markers in two modules, respectively (middle). (D) Left panel: Spatial visualization of REARGCs of two different states in the injured regions of adult axolotl telencephalon from 2 DPI to 20 DPI. Cell state is distinguished by the gene modules in C. Right panel: Spatial visualization of Cell cycle module expression from 2 DPI to 20 DPI. (E) EdU staining indicating the proliferating cells in 10 DPI and 15 DPI. See also Figure S13-S22

Interestingly, we captured a new type of RGC enriched at the edge of wound region, but not in the intact right side of telencephalon sections from 2 to 20 DPI (Figure 3A-B and Figure S15A-B). This injury specific RGCs featured the high expression level of genes, such as *Nes, Krt18, Tnc and Ccna2* (Figure 3A-B and Figure S15C; Table S2), which indicate their stem cell state and high proliferation activity, and we therefore named this RGC type as reactivate RGCs (REARGCs). Strikingly, the number of REARGCs appeared to be significantly increased in the VZ along the medium and lateral pallium region and eventually covered the whole wound area around 20 DPI, and disappeared after 30 DPI (Figure 3B, Figure S13D and Figure S16-S22). Unlike brain wound healing process in mammals, during which microglias and macrophages are the first line of cell types responsible for injury site filling (Li et al., 2021b), REARGCs appeared to be the dominant cell type in the wound area between 2 and 20 DPI, suggesting they may be involved in multiple aspects of tissue regeneration process (Figure 3A-B).

Indeed, the detailed transcriptome examination of REARGCs from 2 to 20 DPI showed two waves of functional module induction. The first is early and transient expression of wound response related genes (Module 1), including *Tgfbi, Runx1, Tnc, Cxcr4* and *Hmox1* at 2 DPI at the wound edge, which were rapidly downregulated thereafter (Figure 3C-D). the second wave featured by continuous induction of proliferation related genes (Module 2) in REARGCs from 10 to 20 DPI when REARGCs and immature NPTXEXs expanded in number (Figure 3C-D and Figure S13D; Table S4). The proliferation feature of REARGCs was further confirmed by EDU staining on sections of 10 DPI and 15 DPI, which showed significant enrichment of EDU positive cells extended from injury site along the VZ region, in comparison with uninjured site (Figure 3E). In contrast, other RGC types retained a moderate and constant expression level of cycling related genes on average throughout regeneration stages. Altogether, our findings here suggest that REARGCs may play dual roles in both early inflammatory response to the wound and then switch to cell propagation to cover the injured region.

### Cellular dynamics from REARGCs to NPTXEXs in injury-induced pallium regeneration

The above results elicit an intriguing hypothesis that REARGCs, but not other RGCs, are the major progenitor cell population that differentiates and restores lost neurons, particularly NPTXEXs. If so, we would expect to capture cell clusters that are between REARGCs and NPTXEXs in terms of differentiation state. To better test the hypothesis and to avoid that our fine sectioning procedure may not capture all types of cells within a single section, we chose to dissect the regenerating telencephalon at 15 DPI along the rostral-caudal axis and made three more consecutive sections from wound center towards the caudal direction in addition to the 15 DPI-1 section shown in Fig 3, including one on the wound edge and two more sections at the closed area for stereo-seq analysis (Figure 4A).

**Figure 4.**
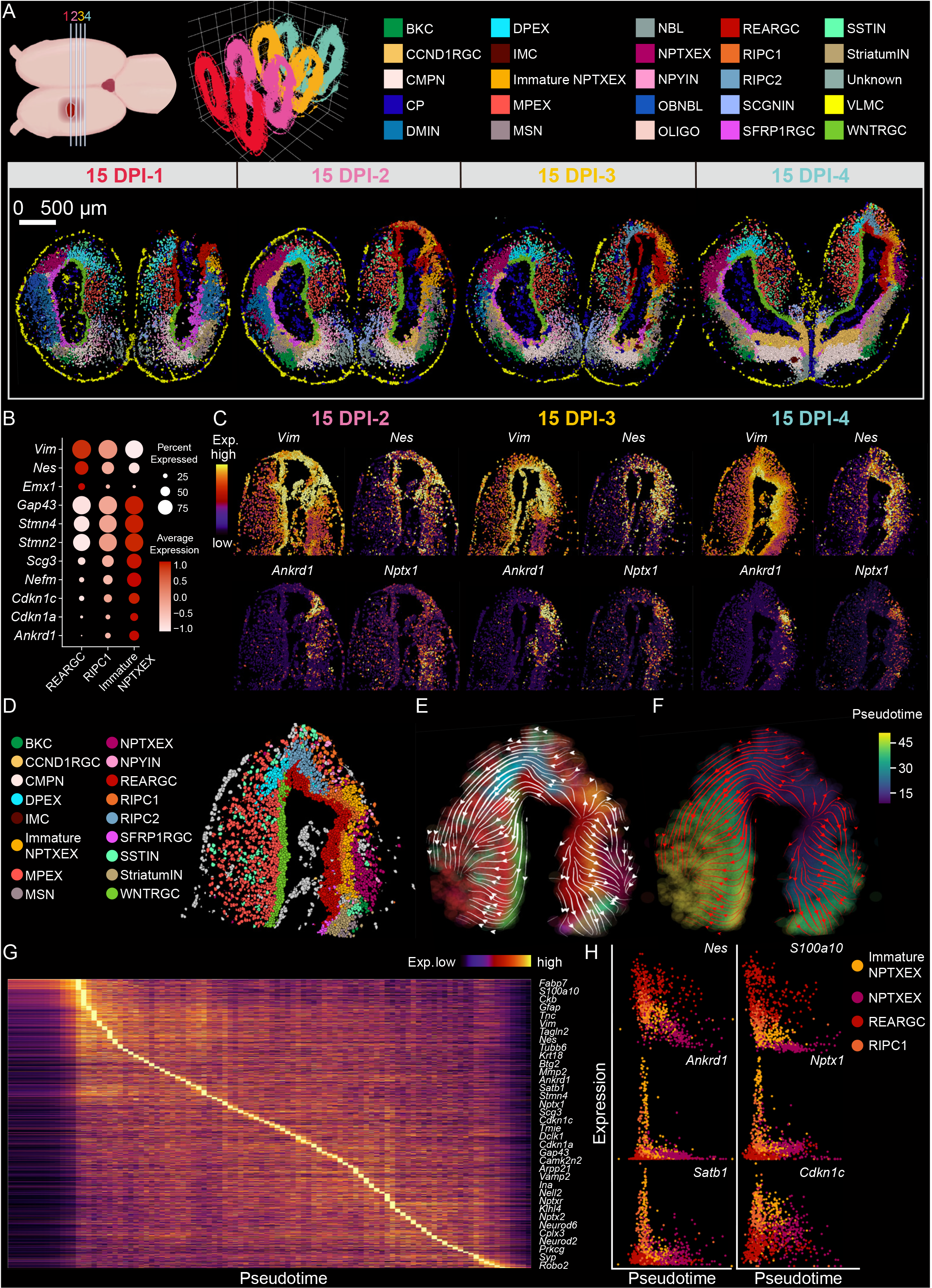
Dynamics of regeneration related cells in differentially recovered sections at 15 DPI reveal potential lineage transition process during regeneration. (A) Positions of 4 continuous sections at 15 DPI (top left) and respective 3D display (top middle). Spatial visualization of cell types identified in 4 continuous axolotl telencephalon sections at 15 DPI at single-cell resolution (bottom). Cell types are annotated by colored cubes on top. Scale bars, 500 μm. (B) Bubble plot reflecting the expression dynamics of marker gene defining REARGCs, RIPC1s and immature NPTXEXs, which are major cell types involved in axolotl telencephalon regeneration. (C) Spatial visualized heatmap showing the expression pattern of key markers for regenerative related cells in the injury area of section 15 DPI-2, 15 DPI-3 and 15 DPI-4. (D) Spatial visualization exhibiting the cell type distribution around the regenerating site in 15 DPI-4. (E-F) RNA velocity streamline plots showing the predicted trajectory of cell lineage transition in the regenerating region of axolotl telencephalon. Areas are colored by either annotated cell clusters (E) or pseudotime (F). (G) Expression heatmap of genes with high transitional activities in a pseudo-temporal order accord with the regeneration process. (H) Scatter plot showing the pseudotime kinetics of *Nes*, *S100a10*, *Ankrd1*, *Nptx1*, *Satb1* and *Cdkn1c* in regenerating cell types. See also Figure S23-S27

By combining data of all four sections, a total of 25 cell types were identified (Figure 4A, Figure S23A-D and S24). NPTXEXs were the major cell population lost at the lesion of the injured hemisphere compared to uninjured lateral side in 15 DPI-1 to 4 sections. REARGCs covered the injury regions in all 4 sections (Figure 3A, Figure 4A and Figure S23D). Interestingly, the NPTXEX population showed a high-to-low spatial gradient from the remote regions in 15 DPI-4 section to the center of the injured area in 15 DPI-1 section (Figure 3A, Figure 4A and Figure S23D), which is consistent with previous micro-CT scanned axolotl regeneration data (Amamoto et al., 2016). These data suggested that the reconstitution of lost neurons probably occurs in accompany with the conjunction of injury edges, the process of which may initiate from the peripheral region towards the center of incision.

We then explored how lost NPTXEXs were restored around the injury site. As expected, the newly formed NPTXEXs in 15 DPI-4 section were nicely located adjacent to immature NPTXEXs and REARGCs, suggesting a possible transition between these cell types (Figure 4A). Excitingly, a regenerative specific cluster of cells were identified in an intermediate state between REARGCs and immature NPTXEXs and therefore named as regeneration intermediate progenitor cells 1 (RIPC1) (Figure 4A; Table S2). They expressed both REARGC markers including *Vim*, *Nes*, *Krt18*, *S100a10* and immature NPTXEX markers including *Ankrd1*, *Stmn4*, *Nptx1* (Figure 4B-C). Expression of cyclin inhibitors *Cdkn1a* and *Cdkn1c* was upregulated in RIPC1 and further upregulated in immature NPTXEXs compared to REARGCs (Dutto et al., 2015; Mademtzoglou et al., 2018), suggesting the proliferation is slowing down along the potential transition axis of REARGC-DIPC1-immature NPTXEX (Figure 4B-C). Thus, our data revealed four nicely adjacent cell layers of REARGCs, RIPC1s, immature NPTXEXs, and mature NPTXEXs, respectively (Figure 4D and Figure S25-S27), inspiring us with a potential linage transition for neurogenesis.

Indeed, cell type based (Figure 4E) and pseudotime based (Figure 4F) RNA velocity analyses on the 15 DPI-4 section also suggested a similar putative lineage transition from REARGCs to RIPC1s, then to immature NPTXEXs and eventually NPTXEXs. Similar observation was made on sections of 15 DPI-2 and −3, too (Figure S28A-C and Figure S28 F-H). To dissect our velocity results in more details, we calculated the genes that show patterned expression change along the pseudotime axis (Figure 4G, Figure S28D and I), which were consistent with their gene function in the putative lineage transition, such like the descending of *Nes* and the ascending of *Cdkn1c* (Figure 4H, Figure S28E and J). In summary, our results imply a potential scenario that REARGCs proliferate to cover the wounding site of injured axolotl telencephalon, then convert or differentiate into RIPCs and eventually produce immature neurons and reconstruct the lost tissue.

### Comparison of the NPTXEX formation processes in development and regeneration

The nicely ordered cell layer distribution and potential lineage transition discovered at injury site are similar to those in the developmental brain, thus prompting us to further compare these two processes. Notably, DRGCs, DIPCs and immature NPTXEXs arrayed from VZ to pallium region were observed at as early as stage 44 (Figure 5B). When the mature NPTXEXs appeared at stage 57, four nicely adjacent cell layers of DRGCs, RIPC1s, immature NPTXEXs, and mature NPTXEXs were observed with high similarity to that in 15 DPI-4 section (Figure 5A-B), indicating a possible recapitulation of NPTXEX development during the injury-induced regeneration. To further test this possibility, all types of RGCs in the dorsal left telencephalon from developmental stage 44, 54 and 57, as well as 15 DPI-4 and control section in injury model were pooled, and applied to correlation analysis. Indeed, the results showed the gene expression pattern of REARGCs is mostly correlated with DRGCs in Stage 57, rather than other RGC types from the same section (Figure 5C). In addition, the spatial expression heatmap of key markers such as *Nes*, *Nptx1* and *Cdkn1c* are much alike between 15 DPI-4 and Stage 57 (Figure 4C, Figure 5D-E, Figure S29 A-C). The RNA velocity analysis has simulated parallel lineage transition trajectories to generate NPTXEXs in developmental and regenerative processes, from RGCs to IPCs to immature neurons to mature neurons (Figure 4E, Figure 5F-H and Figure S29 A-C).

**Figure 5.**
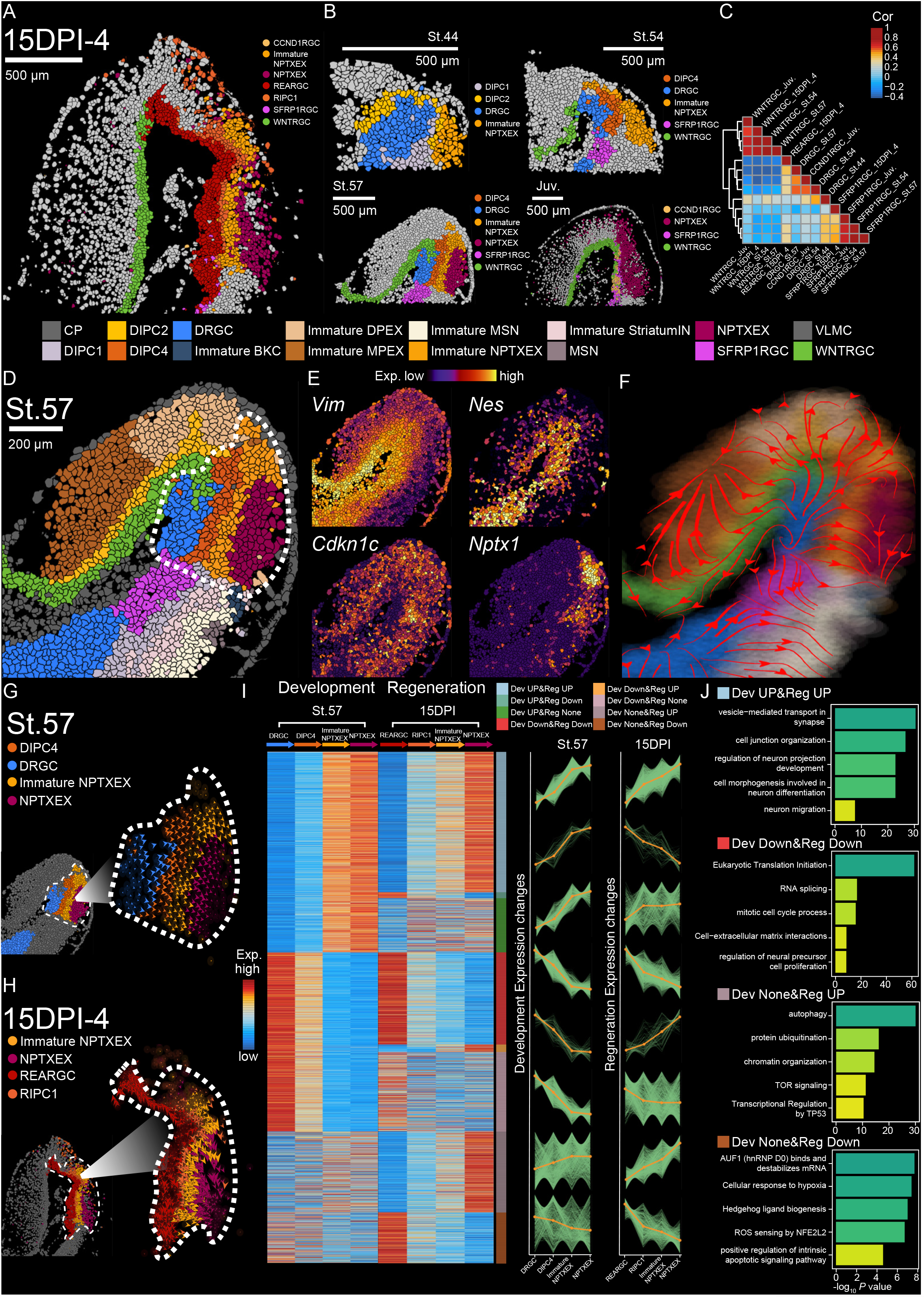
Comparison of neurogenesis in development and regeneration process. (A-B) Spatial distribution of RGCs, IPCs, immature NPTXEXs and NPTXEXs in the dorsal pallium of left hemisphere at 15 DPI-4 (A), and at St.44, St.54, St.57 and Juv. (B). (C) Heatmap represents the correlation between different RGC types across four developmental or regenerative stages. (D) Spatial distribution of dorsal pallium cell types in the left hemisphere of axolotl telencephalon, cells potentially involved in the development of NPTXEX are framed by white dash line. Cell types are annotated by colored cubes on top. (E) Spatial visualized expression of key markers for cells involved in NPTXEX development. (F) RNA velocity streamline plot showing the predicted dorsal pallium development trajectory in the left hemisphere of axolotl telencephalon at St.57. (G-H) RNA velocity streamline plot showing the predicted lineage transition trajectory of RGCs, IPCs, immature NPTXEXs and NPTXEXs at St.57 (G) and 15 DPI-4 (H). (I) Left panel: Heatmap of gene expression during NPTXEX development and regeneration. Eight distinct groups of genes were sorted by their expression dynamic pattern. Right panel: Line plot depicting standardized Stereo-seq signal by fuzzy cluster analysis for eight groups, with green lines representing the expression dynamics of each individual gene and the orange line representing the integrated pattern of each group at Stage 57 and 15 DPI (right). Names of eight groups are listed on top right. Dev UP&Reg UP, co-upregulation in development and regeneration. Dev UP&Reg Down, upregulation in development while downregulation in regeneration. Dev UP&Reg None, upregulation in development while no significant change in regeneration. Dev Down&Reg Down, co-downregulation in development and regeneration. Dev Down&Reg UP, downregulation in development while upregulation in regeneration. Dev Down&Reg None, downregulation in development while no significant change in regeneration. Dev None&Reg UP, no significant change in development while upregulation in regeneration. Dev None&Reg Down, no significant change in development while downregulation in regeneration. (J) Barplot exhibiting the representative GO enrichment pathways of Dev UP&Reg UP, Dev Down&Reg Down, Dev None&Reg UP and Dev None&Reg Down in I. See also Figure S28-S30

We further used potential DRGC to NPTXEX transition at Stage 57 to represent developmental neurogenesis, and potential REARGC to NPTXEX transition of 15 DPI-4 to represent the regenerative neurogenesis, and comprehensively assess the molecular dynamics of these two potential transition processes. We examined the gene expression patterns in the four related cell types along each process, and classified eight groups of gene expression patterns showing similar or opposite trends (Figure 5I; Table S5). Pathway enrichment analysis on these patterned expressing genes revealed that in both development and regeneration, pathways involved in neuronal differentiation, migration, maturation, communication and synaptic activities were up-regulated. However, pathways related to proliferation, cell cycle progression and factors promoting these processes such as translation initiation, RNA splicing were both down-regulated (Figure 5J). These results fit the expected notion that the stemness and proliferation were declined with the onset of neurogenesis (Figure 5J). Interestingly, protein ubiquitination, chromatin organization, mTOR signaling and transcriptional regulation by TP53 were found specifically upregulated in regeneration, suggesting possible immune and metabolism strategies to control the rapid cell growth during regeneration. In addition, we also observed a regeneration specific rise in activities of autophagy, as well as a regeneration specific decline in activities of response to wound and stress (Figure 5J), reflecting a possible transition of molecular machinery from wound response in REARGC to neuron regeneration. Besides the enriched pathways, we also identified hundreds of regenerative specific genes (Table S6). Overall, our data suggested that regeneration to certain extent is the re-initiation of development, but also exhibits its unique features. Though majority of them showed expression pattern as expected, such as *JunD* and *Tnc* that are known to be involved in nervous system regeneration (Chen et al., 2010; Raivich et al., 2004), several unexpected or functionally unknown genes were discovered, such as filament reorganizational *Krt18*, protein phosphorylation related *S100a10*, tumor suppressor *Tagln2* and *Tnfsf10*, and endothelial cell activator *Ankrd1* (Figure S30), which may be interesting targets in following functional studies of brain regeneration.

Altogether, our detailed comparison of spatial transcriptomes revealed great similarity in both spatial distribution and enriched molecular pathway between development and regeneration, suggesting regeneration of axolotl brain may partially recapitulate neurogenesis in brain development through differentiation of stem/progenitor cells with similar molecular regulations.

## DISCUSSION

### Dynamic cell atlas of axolotl telencephalon through development and regeneration at single cell resolution

The goal of regenerative medicine in the brain is to restore not only the intricate tissue architecture and cell composition of the injured region, but also the molecular and cellular homeostasis and functions of recovered brains. While we have learned much about the natural regeneration process by studying various types of animal models, especially the highly regenerative fish and axolotls, key questions including whether the brain of these models can be fully regenerated both anatomically and functionally, how progenitor cells take the regenerative responsibility and how gene activity orchestrates cellular responses upon jury remain unanswered. To this end, a complete set of knowledge regarding the cellular and molecular profiling during development and regeneration in regenerative models is needed in the first place. Though a cellular map of the pallium has been previously built by *in situ* hybridization with a few marker genes in axolotls (Amamoto et al., 2016), more comprehensive cell type identification and their spatial organization and gene activity dynamics in the context of development and regeneration for mechanistic investigation is still lacking.

Taking the advantage of our Stereo-seq and the single nucleus extraction method developed in this study, we successfully established the first single cell level spatial-transcriptomic atlas of axolotl telencephalon, a marked advancement in resolution and throughput in comparison with the brain atlas data in previous studies (Maynard et al., 2021; Ortiz et al., 2020). The single cell level transcriptomics displayed on Stereo-seq sections also empowered us to elucidate the spatial-temporal relationship between diverse cell types in regeneration, between development and regeneration, the knowledge of which is essential but not yet clear. Furthermore, the large size of our stereo-seq chip allowed us to capture the entire telencephalon on the same section, with the left hemisphere injured but the right one uninjured for direct comparison. With these technical advantages, our data provided inspiring information to several important questions in brain regeneration, revealing the key cell populations and their dynamics in the regeneration niche, suggesting a possible strategy for lost tissue reconstitution by lineage-transition from injury-specific RGCs, and elucidating that the injury-induced regeneration partially recapitulate the neuron developmental hierarchy.

### Diversification of RGC subtypes in VZ region through development and identification of transient activated RGCs involved in brain regeneration

RGCs are thought to be ancestor cells that give rise to all cells in brain development and maintained in the VZ region of adult amphibian, featuring by positive staining of BrdU (Joven et al., 2018; Kirkham et al., 2014). Accordingly, four subtypes of RGCs are identified in our study in normal axolotl telencephalon development and maturation. DRGCs that are dominantly distributed along VZ during developmental stages, are shown to be gradually substituted by three locally restricted RGC subtypes (Figure 2A-B), indicating a possible lineage maturation trace of these endogenous neural progenitor cells. Though a decrease of early embryonic markers, the three RGC subtypes in homeostasis retained their feature of NSCs and cell cycle markers, which show similarity to the ependymoglial cells and radial glial cells previously identified in newts and zebrafish, respectively (Lust and Tanaka, 2019). In the red spotted newts, two types of ependymoglial cells, quiescent type I ependymoglial cells and proliferating type II ependymoglial cells, have been reported unevenly distributed along ventricle (Berg et al., 2010; Kirkham et al., 2014). In contrast, we diversified the RGCs in hemostasis based on a combination of whole transcriptome features with their specific localizations, instead of mainly on proliferation status (Figure 2A), thus leading to different classification of RGCs along VZ region. The CCND1RGC, one of the three RGC subtypes in homeostasis stages, is characterized with high cycling gene expression. It is majorly distributed in the VZ region adjacent to the dorsal pallium and to the bed nucleus of the stria terminalis region defined as proliferative hot spots in other salamander species (Kirkham et al., 2014) (Berg et al., 2010), raising the possibility that the CCND1RGCs may share similarity with previously identified type II ependymoglial cells in these spots.

Upon injury, the amphibian brain regeneration is accomplished mainly by resident RGCs/ependymoglial cells through activation and differentiation in response to environmental cues (Berg et al., 2010; Joven et al., 2018; Kirkham et al., 2014), yet how such sequential process is regulated and if other types of cells participate remains largely elusive. By identification of the REARGC and presentation of its dynamic regenerative function, our work not only support the injury-specific appearance of cells with elevated proliferation capacity around the lesion sites (Amamoto et al., 2016; Kirkham et al., 2014), but also indicate that these amplifying RGCs may serve as the cell origin of de novo neurogenesis (Amamoto et al., 2016; Berg et al., 2010; Kirkham et al., 2014).

### Regeneration through REARGCs largely mimics early telencephalon development with DRGC in axolotl

More intriguingly, by comparing our regeneration and development data, it becomes quite clear that REARGCs are in a state similar to DRGCs in terms of gene expression profile and pathway enrichment (Figure 5C and 5J), such as the elevated expression of translation related genes, the typical characteristic of stem cell undergoing active proliferation (Baser et al., 2017). As DRGCs appear from the earliest stage of development we sampled, and presumably give rise to other RGC types and neurons in adulthood, it represents a more primitive type of stem cells with higher multipotency. On the other hand, since DRGCs appear to be consumed in adulthood according to our data, it is mostly likely that REARGCs are originated from resident adult RGCs by reprogramming, though it remains to be determined which type of RGCs are activated and reprogrammed to REARGCs.

As previously suggested, a few possible REARGC origins exist (Amamoto et al., 2016; Berg et al., 2010; Kirkham et al., 2014). The first is the local WNTRGCs or SFRP1RGCs at the lesion site that respond to injury and immediately converted to REARGCs for early wound healing response; The second likely origin is CCND1RGCs, which might be the previously defined proliferative hot spot RGCs responsible for brain neuron maintenance in homeostasis state (Berg et al., 2010; Kirkham et al., 2014). It is possible that REARGCs are directly converted from CCND1RGCs in the VZ region adjacent to the dorsal pallium, where the incision takes place, or originate from ventral CCND1RGCs migrating to the injury site within two days (Figure 3B), the evidence of which requires further investigations. It is of note that at 10 and 15 DPI, more RGCs at the wound area harbor high EdU labeling activity than that at the VZ region (Figure 3E), suggesting REARGCs may locally replenish its own population, consistent with previous reports (Amamoto et al., 2016). In any case, future VZ-region specific labeling or functional perturbation assay are required to eventually elucidate how RGCs at different regions function during regeneration.

### Comparison of molecular and cellular features between axolotl brain regeneration and mammalian brain injury recovery

Inflammation/immune-responses have been reported to be critical to lead to a successful regeneration (Kyritsis et al., 2012). Unlike the axolotl, lesion in mammalian adult brains often leads to tissue loss and cystic scar formation, with very limited functional recovery (Hagberg et al., 2012). Only low-cycling/quiescent NSCs are identified in defined areas of postnatal mammalian brains, and they are generally difficult to be activated to give rise to proper cell types to repair brain lesion (Furutachi et al., 2015; Llorens-Bobadilla et al., 2015; Yang et al., 2007). The pro-inflammatory molecules secreted at the lesion following hypoxia insult are thought to be detrimental to neurogenesis in mammals, and such negative effects are elevated by the long-term persistence of glial scar (Silver and Miller, 2004). Interestingly, a transit wave of wound-response is also observed in REARGCs at the edge of the lesion site during cortex regeneration, which represented by the upregulation of inflammation and hypoxia related genes at 2 and 5 DPI (Figure 3C-D). It suggests a similar early response in axolotl as those in mammalian system. In contrast to the sustained inflammation in injured mammalian brain, REARGCs in the injured axolotl pallium decrease their activity in wound-response and transform their molecular features into proliferation around 10 DPI (Figure 3C-E), raising the possibility that they may start to function as NSCs to initiate de novo neurogenesis, the process of which is further proved by the nicely arranged cell layers re-exhibiting in the developmental process (Figure 5). Based on these observations, we predicted that axolotl REARGCs may play dual roles in injury-induced brain regeneration: firstly, respond to early wound insults and then expand to reconstitute lost neurons. The injury-specific REARGC population and the well-controlled transition between two functions may provide key mechanism to balance inflammation versus neurogenesis, which endows axolotl the capacity of neuronal regeneration.

In summary, our work provides the most comprehensive atlas of the axolotl telencephalon to date. Such large-scale efforts not only prompt to redefine the subpopulation of radial glial cells based on whole transcriptome feature, but also start to reveal their dynamic transition and functions in development and regeneration with significant similarity at molecular level. In the future, advanced technology with higher resolution and RNA capture capability, as well as accurate cell membrane border definition strategies, will enable the identification of cell types and transcript features in a more accurate way. Moreover, as injury-induced regeneration would be joint behaviors in different regions, including cells from the olfactory bulb, yet limitation may exist for full recovery (Maden et al., 2013) (Amamoto et al., 2016). Therefore, it could be of great interest in the future study to perform continues sections that enable 3D reconstruction of multiple brain regions in a longer regeneration time, which would help display the networks of neuron projections and connections during regeneration and to investigate whether they are rebuild completely. 3D map can also let us investigate whether molecular and cellular cues of regeneration are polarized along rostro-caudal axis.

## ACKNOWLEDGMENTS

This work was supported by the Guangdong Provincial Key Laboratory of Genome Read and Write (2017B030301011), the National Natural Science Foundation of China (32171289, 31970619) and the Innovative Research Group Program of Hubei Province (2020CFA017). Ji-Feng Fei was supported by the National Key R&D Program of China (2019YFE0106700), the Natural Science Foundation of China (31970782), Project of Department of Education of Guangdong Province (2018KZDXM027), Key-Area Research and Development Program of Guangdong Province (2018B030332001, 2019B030335001) and Guangdong-Hong Kong-Macao-Joint Laboratory Program (2019B121205005).

## AUTHOR CONTRIBUTIONS

X.W., G.Y., J.F.F., X.X., C.L. and H.L. conceived the idea; G.Y., J.F.F., X.X., C.L. and H.L. supervised the work; X.W., S.F., H.L. and J.F.F. designed the experiment; S.F., X.L. and N.Z. performed the majority of the experiments with the help from M.C., J.J., J.X., Y.Z., P.L., X.S. and C.P.; X.W., Y.L., S.W., W.F. and Y.Y. performed data analysis; M.K. and T.Y. performed database construction; Y.L., X.Q., L.W., L.H., L.C., Y.Y., X.P., S.G., A.C., M.A.E., H.Y., J.W., G.F. and L.L. gave the relevant advice; H.L., Y.G., L.C., J.F.F. and X.W. wrote the manuscript with input from all authors. All other authors contributed to the work. All authors read and approved the manuscript for submission.

## RESOURCE AVAILABILITY

### Lead contact

Further information and requests for the resources and reagents may be directed to the corresponding author Ying Gu

### Material availability

All materials used for Stereo-seq are commercially available.

### Data and code availability

All raw data generated by Stereo-seq have been deposited to CNGB Nucleotide Sequence Archive (accession code: CNP0002068 (https://db.cngb.org/search/project/CNP0002068). Any additional information is available from the corresponding authors upon reasonable request.

## METHODS

### Animal care

The d/d Strain of *Ambystoma mexicanum* used in this study was originally obtained from Elly M. Tanaka laboratory (Research Institute of Molecular Pathology, Vienna Biocenter, Vienna, Austria). Animals were housed and bred at 18-20°C in fresh water under standard conditions. All relevant procedures of animal experiments were carried out in accordance with the animal welfare legislation in China, with local approval from the Biomedical Research Ethics Committee of Guangdong Provincial People’s Hospital.

### Brain injury

10-13cm juvenile axolotls were used for brain injury experiments. Axolotl brain injury was performed as described previously (Amamoto et al., 2016). Briefly, animals were firstly anesthetized in 0.03% ethyl-p-aminobenzoate solution (E1501, Sigma-Aldrich, St.Louis, MO), followed by surgeries to create rectangular cranial skin/skull flaps and expose the left telencephalon of each experimental animal using scalpels and spring scissors. Finally, a square-shaped (size 0.5mm x 0.5mm) piece of dorsal telencephalon tissue was removed for each animal to generate brain damage. To accurately determine the injury site, the incisions were placed right in between the olfactory bulb and choroid plexus on the left telencephalon of animals. After the injury, the cranial skin/skull flaps were restored without suture.

### Tissue collection

For stereo-seq cryosection, brain samples from three developmental stages (Stage 44, stage 54, stage 57), juvenile, adult and metamorphosed animals, and seven regenerative stages (2 days post injury (2 DPI), 5 DPI, 10 DPI, 15 DPI, 20 DPI, 30 DPI, 60 DPI) were collected from ethyl-p-aminobenzoate anaesthetized d/d axolotls. Brain samples were immediately snap-frozen in Tissue-Tek OCT (4583, Sakura, Torrance, CA) with liquid nitrogen prechilled isopentane and then transferred to −80°C refrigerator for storage before further operation. To minimize RNA degradation, the entire dissection procedure was performed on ice, and the tissue collection was completed within 30 minutes.

For *in situ* hybridization and EdU detection, we collected additional brains from juvenile and adult d/d axolotls for cryosection. Brain samples were first fixed with MEMFA for 3 days, then transferred into 30% sucrose prepared in 1× PBS for 24 hours and finally embedded in Tissue-Tek OCT with dry-ice. The OCT blocks were stored in −80°C freezer before cryosection.

### Tissue cryosection

For cryosection collection, the working area of the freezing microtome (CM1950, Leica, Wetzlar, Germany) were sequentially cleaned with RNase Zap (AM9780, Invitrogen, Waltham, MA) and DPEC (40718, Sigma-Aldrich, St. Louis, MO)-treated water. After the machine is completely dried, we set the machine temperature to −25°C for cryosection. We collected 20-μm coronal cryosections for Stereo-seq, and 10-μm coronal cryosections for *in situ* hybridization and EdU detection, according to manufactory’s instructions.

### Stereo-seq tissue fixation and ssDNA staining

For Stereo-seq cryosection collection, tissue sections were directly adhered to the Stereo-seq chip surface. The sections on chips were incubated at 37°C for 3 minutes, followed by methanol fixation for 30 minutes at −20°C. Then the sections on chips were stained with nucleic acid dye (Thermo fisher, Q10212) for ssDNA visualization. Images of ssDNA were acquired with a Ti-7 Nikon Eclipse microscope.

### Stereo-seq libraries construction

Tissue sections placed on the chip were permeabilized using 0.1% pepsin (Sigma, P7000) in 0.01 M HCl buffer and incubated at 37°C for 12 minutes. RNA released from the permeabilized tissue and captured by the DNA nanoball (DNB) was reverse transcribed at 42°C overnight. Tissue sections were digested with tissue removal buffer (10 mM Tris-HCl, 25 mM EDTA, 100 mM NaCl, 0.5% SDS) at 37°C for 30 minutes after reverse transcription. cDNA-containing chips were then subjected to Exonuclease I (NEB, M0293L) treatment for 1 hour at 37°C and cDNAs were amplified with KAPA HiFi Hotstart Ready Mix (Roche, KK2602). PCR reactions were conducted as follow: 95°C for 5 minutes, 15 cycles at 98°C for 20 seconds, 58°C for 20 seconds, 72°C for 3 minutes and a final incubation at 72°C for 5 minutes. The concentrations of the resulting PCR products were quantified by Qubit™ dsDNA Assay Kit (Thermo, Q32854). A total of 20 ng of DNA were then fragmented with in-house Tn5 transposase at 55°C for 10 minutes, after which the reactions were stopped by the addition of 0.02% SDS. Fragmentation products were amplified with KAPA HiFi Hotstart Ready Mix. The reaction flow was: 1 cycle of 95°C 5 minutes, 13 cycles of (98°C 20 seconds, 58°C 20 seconds and 72°C 30 seconds) and 1 cycle of 72°C 5 minutes. PCR products were purified and used to generate DNB, then sequenced (35 bp for read 1, 100 bp for read 2) on a MGI DNBSEQ-T1 sequencer.

### *In situ* hybridization

Total cDNA was prepared using the RNA mixture from axolotl brain tissues at different stages. Target genes were amplified with total cDNA as template, and synthesized oligonucleotides harboring T7 promoter as primers. The PCR products were used as template to synthesize Dig-labelled antisense RNA probes by *in vitro* transcription. The *in situ* hybridization was performed on 10-μm axolotl brain coronal cryosection as previously described (*Knapp et al., 2013*). Briefly, air-dried slides were washed by 0.1% Tween in 1 × PBS for 5 minutes for three times, then by 0.3% Triton in 1 × PBS for 20 minutes at room temperature. For pre-hybridization, slides were immersed in hybridization buffer (10% dextran, 5 × SSC, 50% formamide, 0.1% Tween, 1 mg/ml yeast RNA, 100 μg/ml heparin, 1 x Denhardt’s solution, 0.1% CHAPS and 5 mM EDTA) at 60°C for at least 30 minutes. Then, slides were incubated in hybridization buffer containing 500 ng/ml antisense RNA probes at 60°C overnight. After hybridization, excessive RNA probes were removed by washing twice with post-hybridization buffer (5 × SSC, 50% formamide and 0.1% Tween) for 30 minutes at 60°C twice, followed by wash buffer (2 × SSC, 50% formamide and 0.1% Tween) for 30 minutes at 60°C. The slides were then washed with 0.2 × SSC buffer (0.2 × SSC and 0.1% Tween) for 30 minutes at 60°C once and room temperature once, followed by TNE buffer (10mM Tris pH7.5, 500 mM NaCl, 1 mM EDTA) for 10 minutes twice. Next, RNase A treatment was performed (Sigma, R4642) in TNE buffer for 1 hour at 37°C. RNase A was removed by washing with TNE buffer for 10 minutes twice. For sample blocking, slides were incubated with MAB buffer (100 mM maleic acid, 150 mM NaCl, 0.1% Tween and pH was adjusted to pH 7.5 with NaOH) for 5 minutes twice and 20 minutes once, followed by incubation with blocking buffer (Roche, 11096176001) at room temperature for 1 hour. Then sections were incubated with Anti-DIG-AP fab antibody (Roche, 11093274910) prepared in blocking buffer overnight at 4°C. Afterwards, slides were washed five times with MAB buffer for 10 minutes at room temperature and once in freshly-prepared AP buffer (100 mM Tris pH 9.5, 50 mM MgCl_2_, 100 mM NaCl and 0.1% Tween). Finally, we used BM purple (Roche, 11442074001) substrate for signal visualization and the reaction was stopped with 1 mM EDTA in 1 × PBS. Primers used for antisense RNA probe synthesis are listed in Table S7.

### EdU labeling

For *in vivo* cell proliferation detection during brain regeneration, a single pulse of EDU (A10044, Thermo Fisher Scientific, Waltham, MA, 10 μg EdU per gram of body weight) was intraperitoneally injected into animals 6 hours prior to sample collection. EdU detection was performed according to the manufacturer’s protocol of the Click iT Plus EdU Alexa fluor 594 imaging kit (C10339, Thermo Fisher Scientific, Waltham, MA).

### Stereo-seq raw data processing

Stereo-seq fastq files were generated from a MGI DNBSEQ-T1 sequencer. Read 1 contained CID and MID sequences (CID: 1-25bp, MID: 26-35bp), while read 2 contained the cDNA sequence. Retained reads were then aligned to the reference genome AmexG_v6.0 (Schloissnig et al., 2021) via STAR (Dobin et al., 2013). Mapped reads with MAPQ ≥ 10 were annotated, then calculated by handleBam (available at https://github.com/BGIResearch/handleBam). Reads overlapped more than 50% with the exon region were counted as exon transcripts. Reads overlapped less than 50% with the exon region yet possess overlapped sequences with the adjacent intron sequence were annotated as intron transcripts, otherwise as intergenic transcripts. UMIs with both the same CID and gene locus was collapsed, and 1 base pair of mismatches to correct sequencing was allowed for PCR errors. Finally, the exonic reads were used to generate a CID-containing expression profile matrix.

### Circling method for single cell identification of Stereo-seq data

The ssDNA images were used to identify the nuclei region via scikit-image python package (version 0.18.1) (van der Walt et al., 2014). Global threshold was applied to filter background noise and Gaussian-weighted local threshold was calculated to process gray images into binary images, with block size of 41 and offset of 0.03. To segment cell nuclei with overlapped regions, the exact Euclidean distance transformation was performed. Labels representing different cell nuclei were transferred to pinpoint DNBs corresponding to spatial positions by watershed algorithm (Roerdink and Meijster, 2000).

### Unsupervised spatially-constrained clustering of Stereo-seq data

The raw count matrices of axolotl adult telencephalon samples were normalized by SCTransform function in Seurat (v4.0.2) (Hao et al., 2021), and spatial information was taken into consideration for unsupervised clustering by custom script. In brief, the mean values of x-axis and y-axis on bins of each single-nuclei cell were calculated as single-nuclei cell space coordinates. Spatial k-nearest neighbor graph 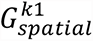 (*k*_1_ = 6) was built via Squidpy and then took the union with the k-nearest neighbor graph 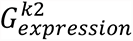 based on transcriptomic data (*k*_2_ is by default set to be 30). The combined graph 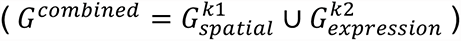 was then used as input for leiden clustering at resolution of 0.5. The adult telencephalon brain regions were identified based on anatomical structure (González et al., 2017; Lazcano et al., 2021).

### Single cell level clustering and annotation of Stereo-seq

Reads captured by DNBs were summarized based on the results of cell segmentation to obtain the single cell level gene expression matrices for downstream analysis. Nuclei with UMIs number less than 200 were discarded. The raw count matrices were then normalized by SCTransform function in Seurat (v4.0.2) to eliminate sequencing depth effects. Reference-based integration was applied along with reciprocal PCA (RPCA) to integrate all slides to avoid potential batch effects. Cluster analysis was performed by FindClusters function in Seurat (v4.0.2) with resolution of 2. Clustering results were displayed by uniform manifold approximation and projection (UMAP) dimension reduction analysis. Major cell types were annotated based on marker gene sets. Visualization of spatial plots in single-DNB resolution with cell boundary were performed by custom script.

### Gene Module analysis

Gene module expression score was calculated to evaluate the expression level of individual cells regarding predefined gene sets. Stemness, cell cycle *and* translation scores of each cell were calculated by AddModuleScore function with default parameters (ctrl = 100, nbin = 24) implemented in the Seurat (v4.0.2) R package (Hao et al., 2021), with respective gene sets manually selected from GO, BIOCARTA, PID, REACTOME and KEGG databases. The detailed gene lists can be found in Table S3.

### Construction of one-to-one orthologous between axolotl and mouse

To optimize the gene annotation of axolotl, a gene ortholog table was created with the mouse genome as the reference gene list. 46,581 axolotl protein coding genes were identified from the AmexG_v6.0 genome (Schloissnig et al., 2021). Mouse reference genome and GFF file (GRCm39) were downloaded from NCBI, the longest transcript of each mouse gene was picked by an in-house Perl script. From which, 22,173 mouse protein coding genes were identified. The orthologous genes were inferred using blastp (Altschul et al., 1997) of the axolotl proteins vs. the mouse proteins. As a result, 20,650 one-to-one orthologous genes were identified between axolotl and mouse, which were used for functional annotation of axolotl in the follow-up analysis.

### Mouse data processing

To compare the expression of stemness related genes in developmental stages between axolotl and mouse, we used the published matrix data and metadata of mammalian cerebral cortex atlas from the Single Cell Portal (https://singlecell.broadinstitute.org/single_cell/study/SCP1290/molecular-logic-of-cellular-diversification-in-the-mammalian-cerebral-cortex) (Di Bella et al., 2021). The count matrices were then normalized by SCTransform function in Seurat (v4.0.2) to eliminate sequencing depth effects. Highly variable genes within the homeotic genes in both axolotl and mouse were identified by SelectIntegrationFeatures function of Seurat (v4.0.2). Based on that, axolotl development data and the mouse cerebral cortex atlas were integrated by FindAnchors and IntegrateData functions of Seurat (v4.0.2) with default dimensionality of 30.

### Mfuzz analysis

Stereo-seq SCT normalized gene expression for cells from the same stage was aggregated to form a pseudo-bulk gene expression matrix. Prior to clustering, genes with expression < 0.5 at all stages were removed. The normalized data of the remaining genes was then Z-score transformed before executing the c-means fuzzy clustering the time-course regeneration data, with two centers and a cluster membership threshold of 0.8 (Kumar and M, 2007).

### RNA velocity analysis

RNA velocity was performed by Dynamo following the instructions at https://dynamo-release.readthedocs.io/ (Qiu et al., 2021). The relative abundance of nascent (unspliced) and mature (spliced) mRNA can be exploited to estimate the rates of gene splicing and degradation. The unspliced and spliced raw count matrices were calculated by Velocity according to annotation file and bam file processed and annotated by handleBam (bin-point label was substituted by cell ID) (La Manno et al., 2018). Unspliced and spliced raw count matrix of regeneration-related cell types were extracted to process by *recipe_monocle* function in Dynamo (Qiu et al., 2021). Highly expressed genes with significant MoranI index were selected as feature genes to perform dimension reduction via UMAP with default parameters. Then, kinetic parameters and gene-wise RNA velocity vectors were estimated on the normalized matrix, which were projected into the visualized spatial plot to retain spatial information. Streamlines were used to visualize the velocity vector flows on specific regeneration-related cell types in injured dorsal region of telencephalic sections.

To facilitate the understanding of related genes determining the fate of target cells, the continuous vector field in the UMAP space and spatial space were established by the vf VectorField function depending on sparseVFC to learn the high dimensional vector field in the expression space from sparse single nuclei cell velocity vector robustly. Next, based on the learned vf VectorField function, ddhodge function was applied on our data to obtain UMAP pseudotime and spatial pseudotime. Finally, the gene expression dynamics vector field was visualized along the pseudotime, and genes with significant spatio-temporal expressional preference were selected.

---

**Figure S1.**
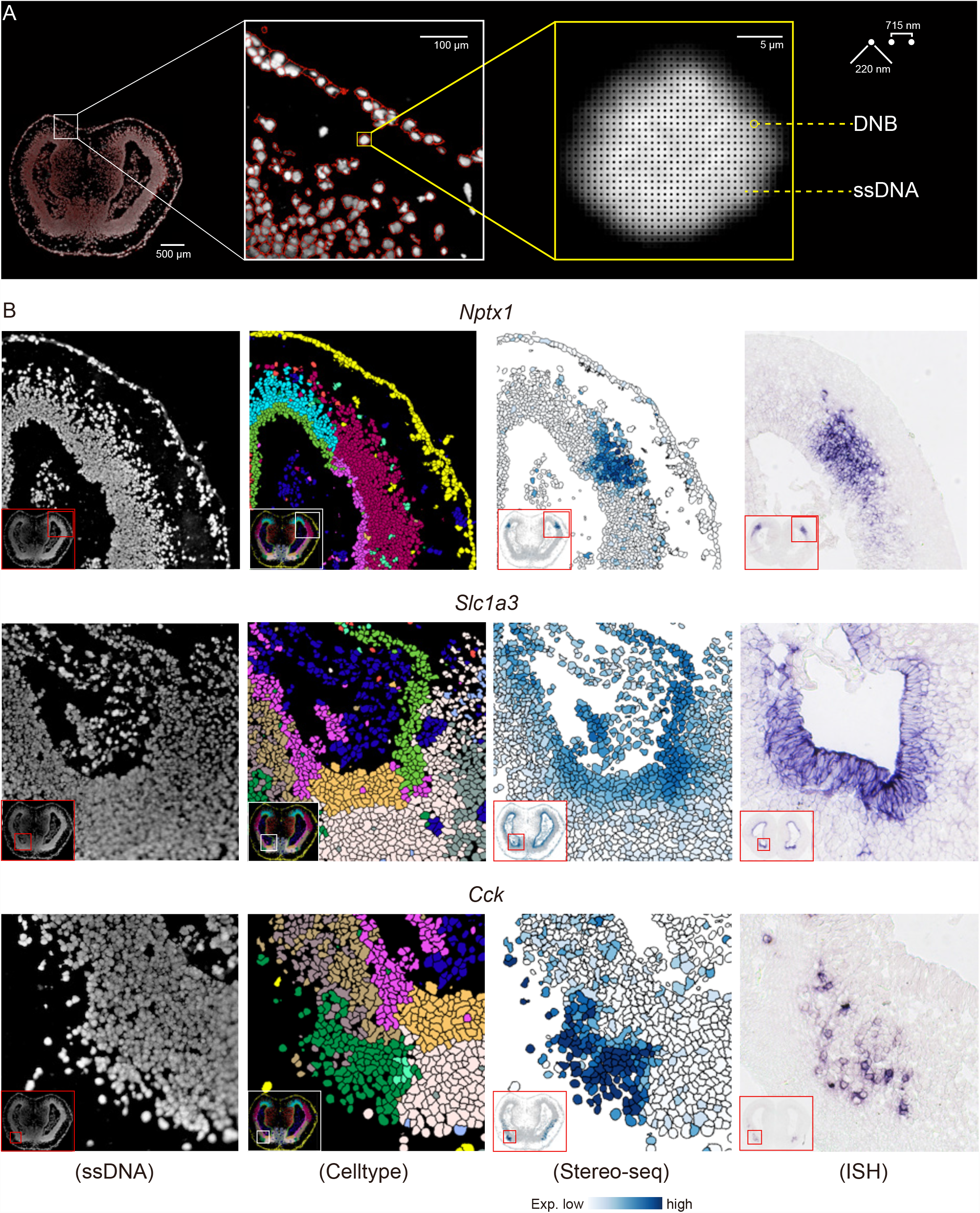
Stereo-seq at single-cell resolution, related to Figure 1. (A) The arrangement of DNB array used to capture mRNA in single nucleus. (B) Images of ssDNA staining, spatial visualization of cell types, spatial visualization of gene expression and corresponding *in situ* hybridization of *Nptx1*, *Slc1a3* and *Cck*.

**Figure S2.**
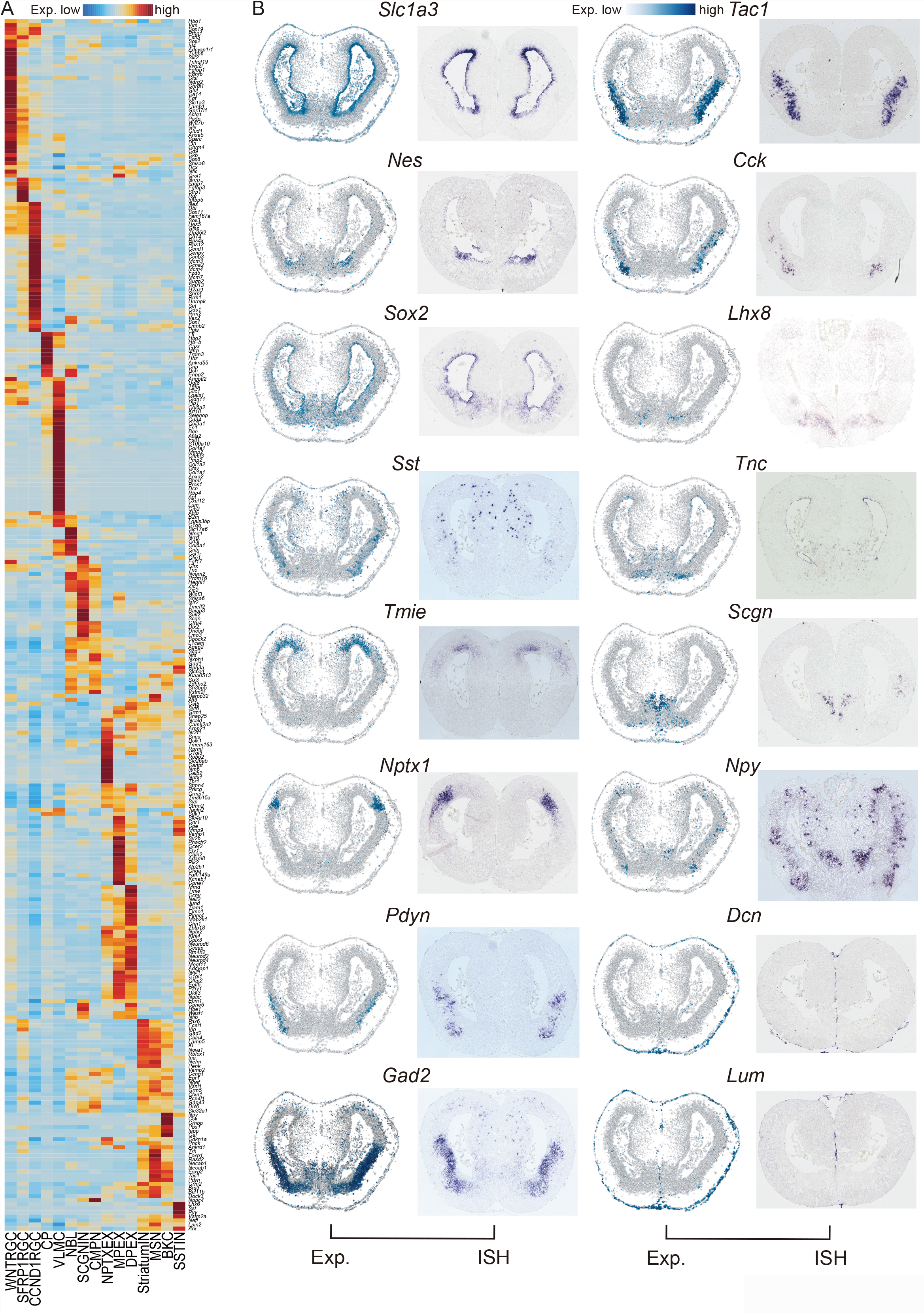
Spatial visualization of marker genes in the adult axolotl telencephalon, related to Figure 1. (A) Heatmap showing the mean expression of DEGs between the 15 cell types in Figure 1D and E. (B) Spatial visualization of representative cell type markers in adult axolotl telencephalon section at single-cell resolution (left). *In situ* hybridization of representative cell type markers in axolotl telencephalon section (right).

**Figure S3.**
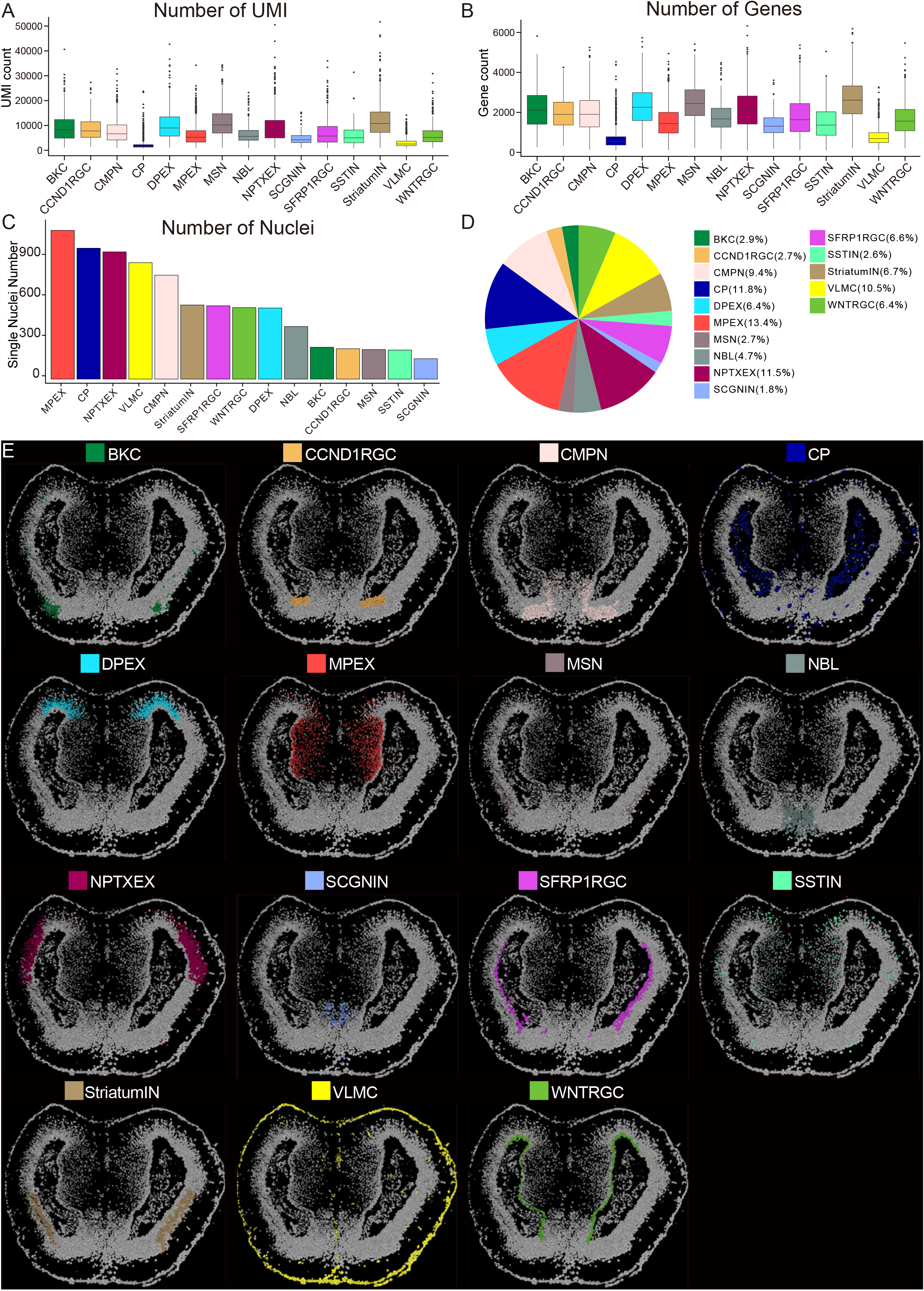
Cell type distribution of adult axolotl telencephalon, related to Figure 1. (A-B) Boxplot showing the count number of UMIs (transcripts) (A) and genes (B) captured by Stereo-seq in 15 cell types of adult axolotl telencephalon, related to Figure 1D and E. (C) Barplot exhibiting the number of single-nucleus for each cell type analyzed in Figure 1. (D) Pie chart reflecting the ratio of each cell type detected in adult axolotl telencephalon. (E) Spatial distribution of each cell type on adult axolotl telencephalon section, cell types are colored as annotated in Figure 1.

**Figure S4.**
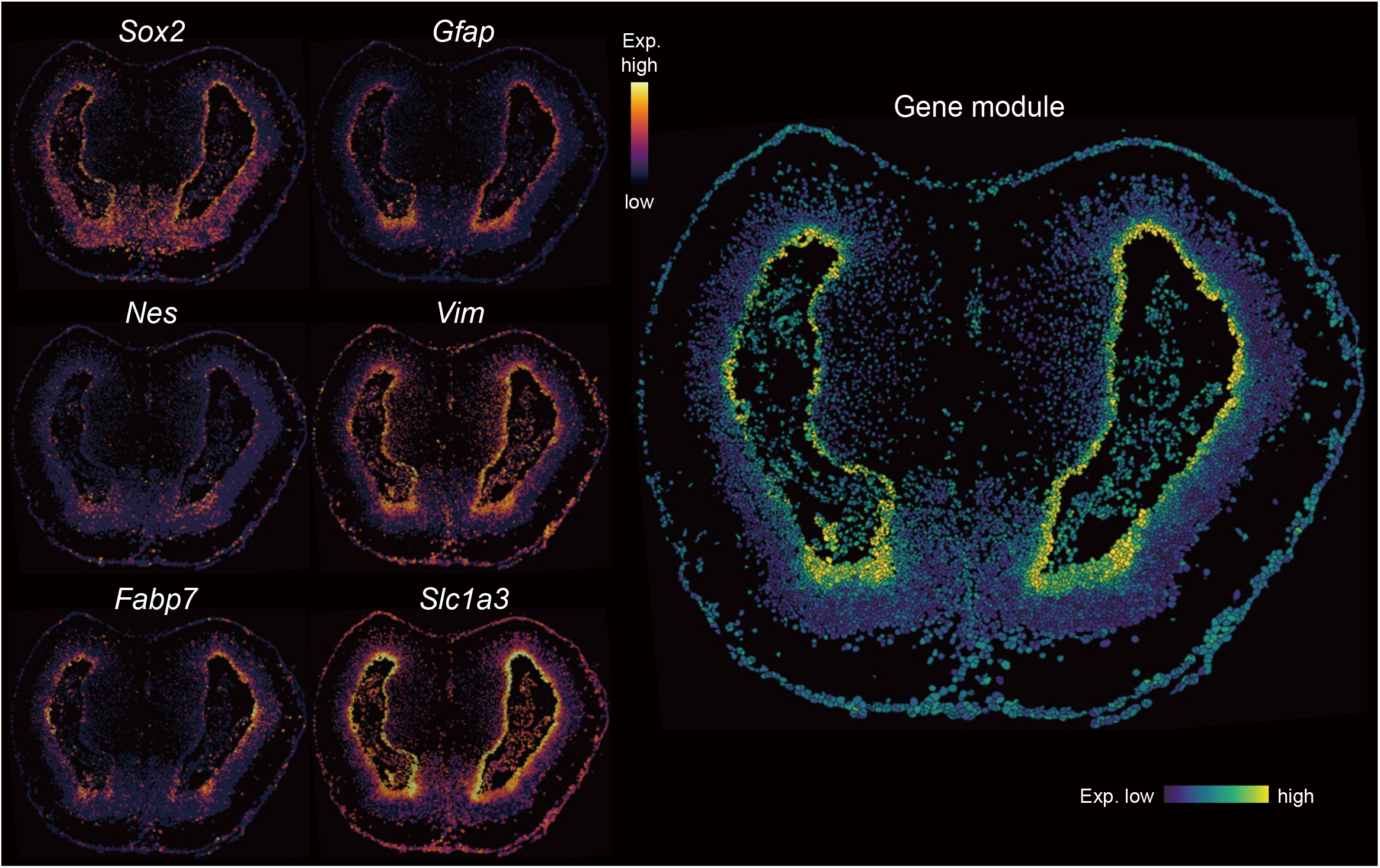
Stereo-seq identifies ventricular zone (VZ) cells in the adult axolotl telencephalon, related to Figure 1. Spatial visualization of radial glial marker expression in VZ on adult axolotl telencephalon section. Expression of markers are exhibited at single-cell resolution (left). Integrated expression of 6 markers is displayed as unified gene module (right).

**Figure S5.**
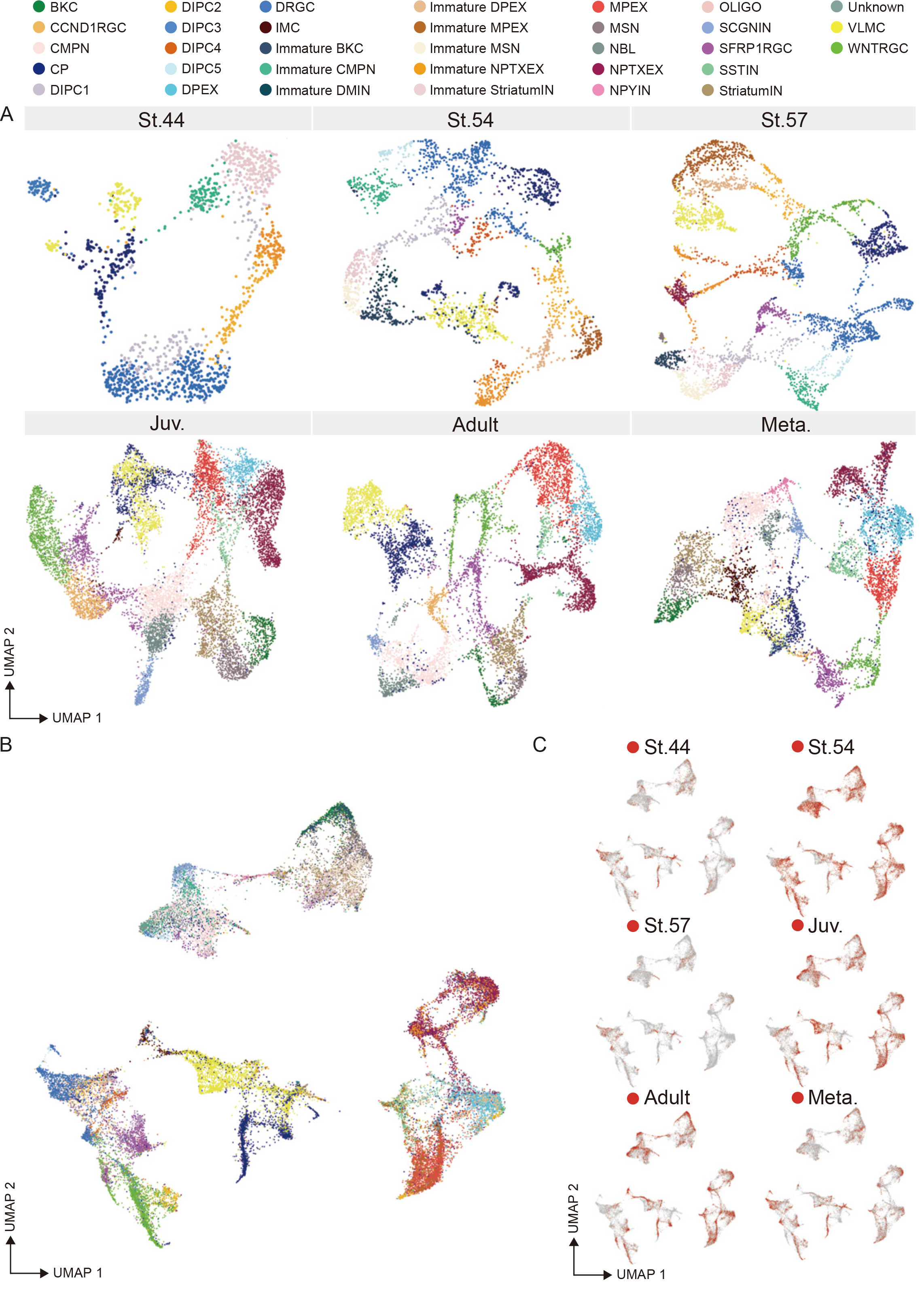
Unsupervised clustering of cells in axolotl telencephalon development, related to Figure 2. (A) UMAP showing the clusters in each stage in Figure 2. 33 cell types are identified and annotated as in Figure 2A. (B) UMAP showing the integration of 33 cell types across different stages. (C) UMAP showing the distribution of cells from each sampling stage in Figure 2. Red dots represent the cells from the corresponding time point, gray dots represent all the other cells.

**Figure S6.**
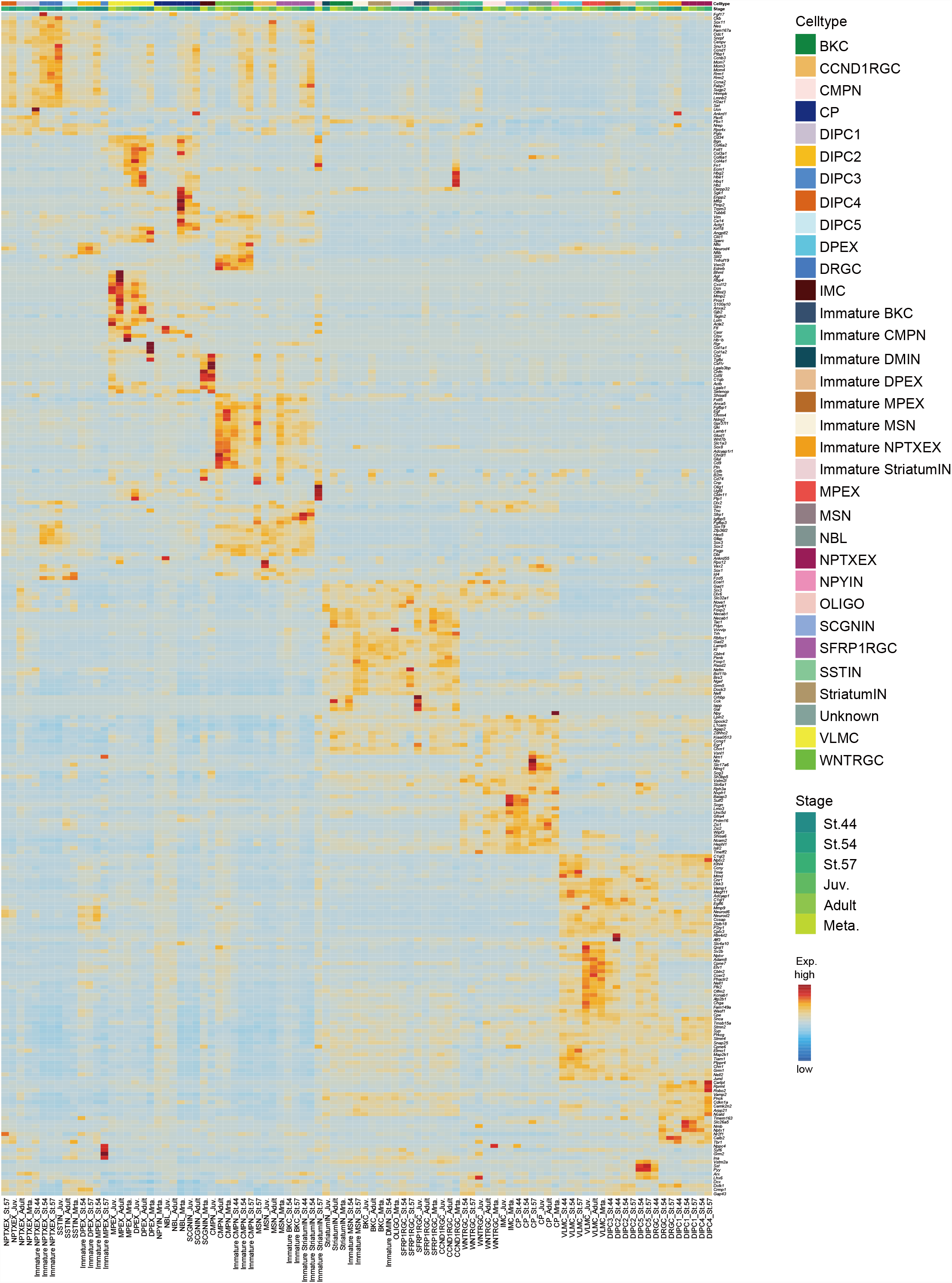
Markers of cell cluster in axolotl telencephalon, related to Figure 2. Heatmap showing the normalized expression of marker genes for the indicated anatomical structures of the axolotl telencephalon sections profiled by Stereo-seq in Figure 2A.

**Figure S7, S8, S9, S10, S11.**
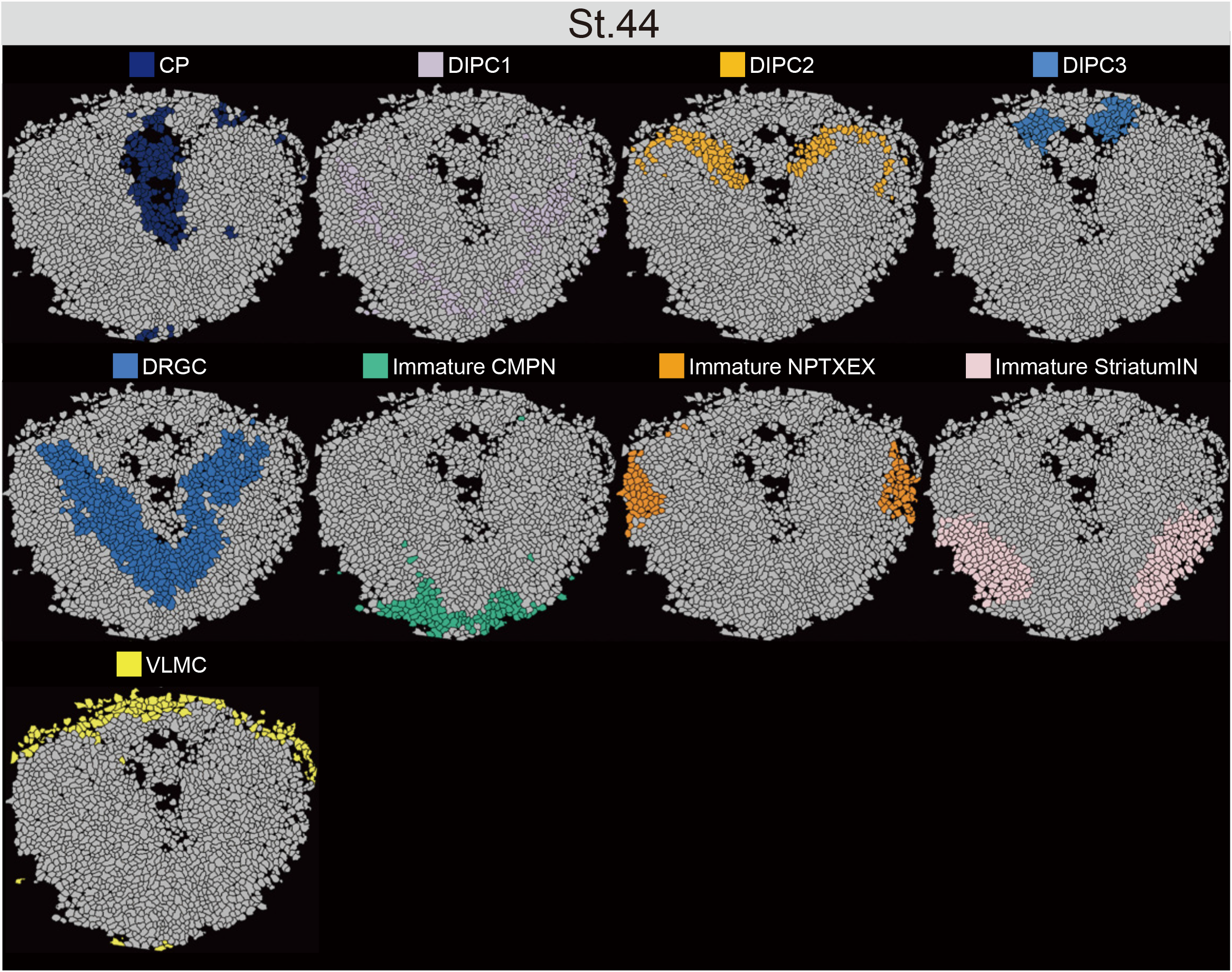
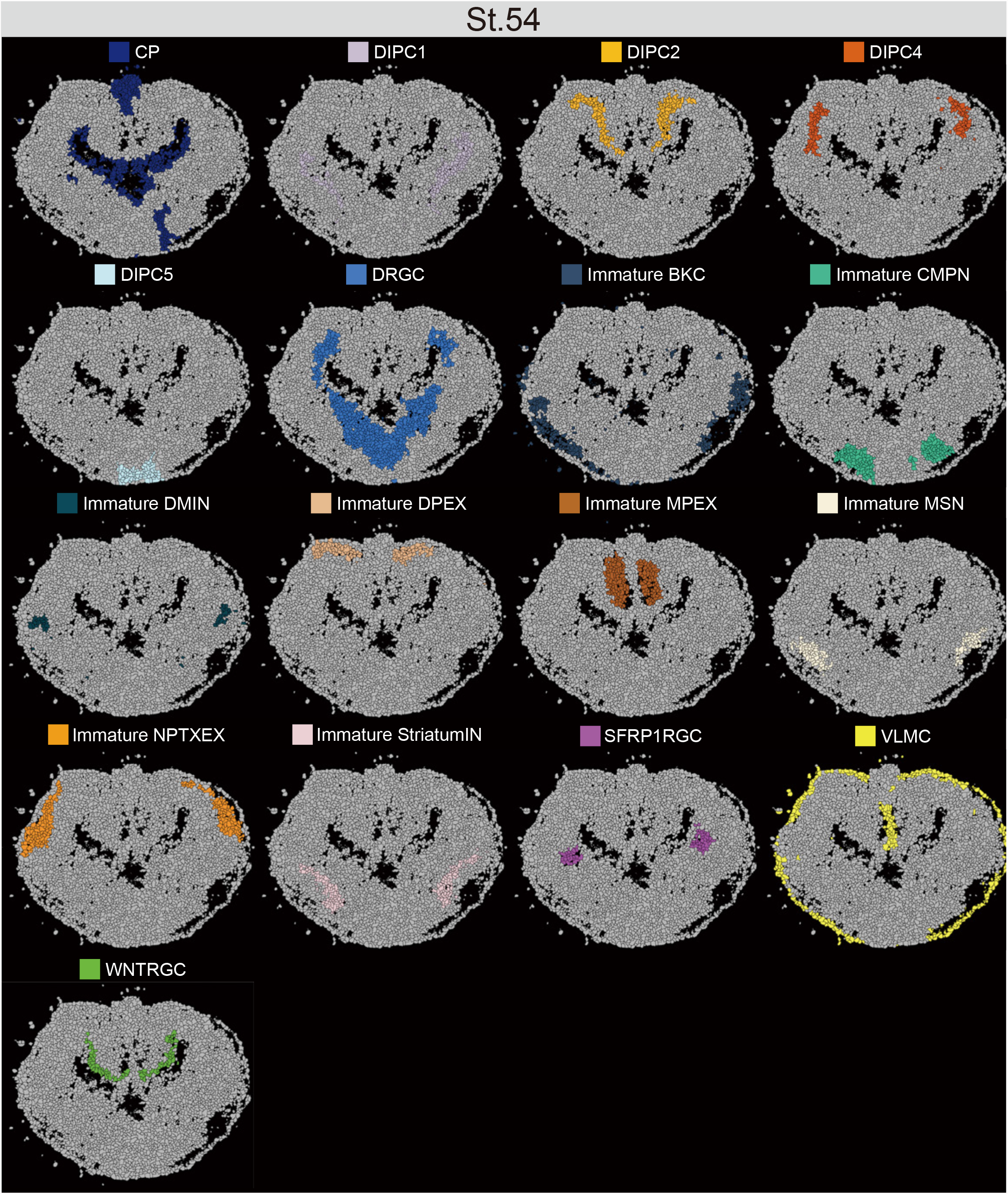
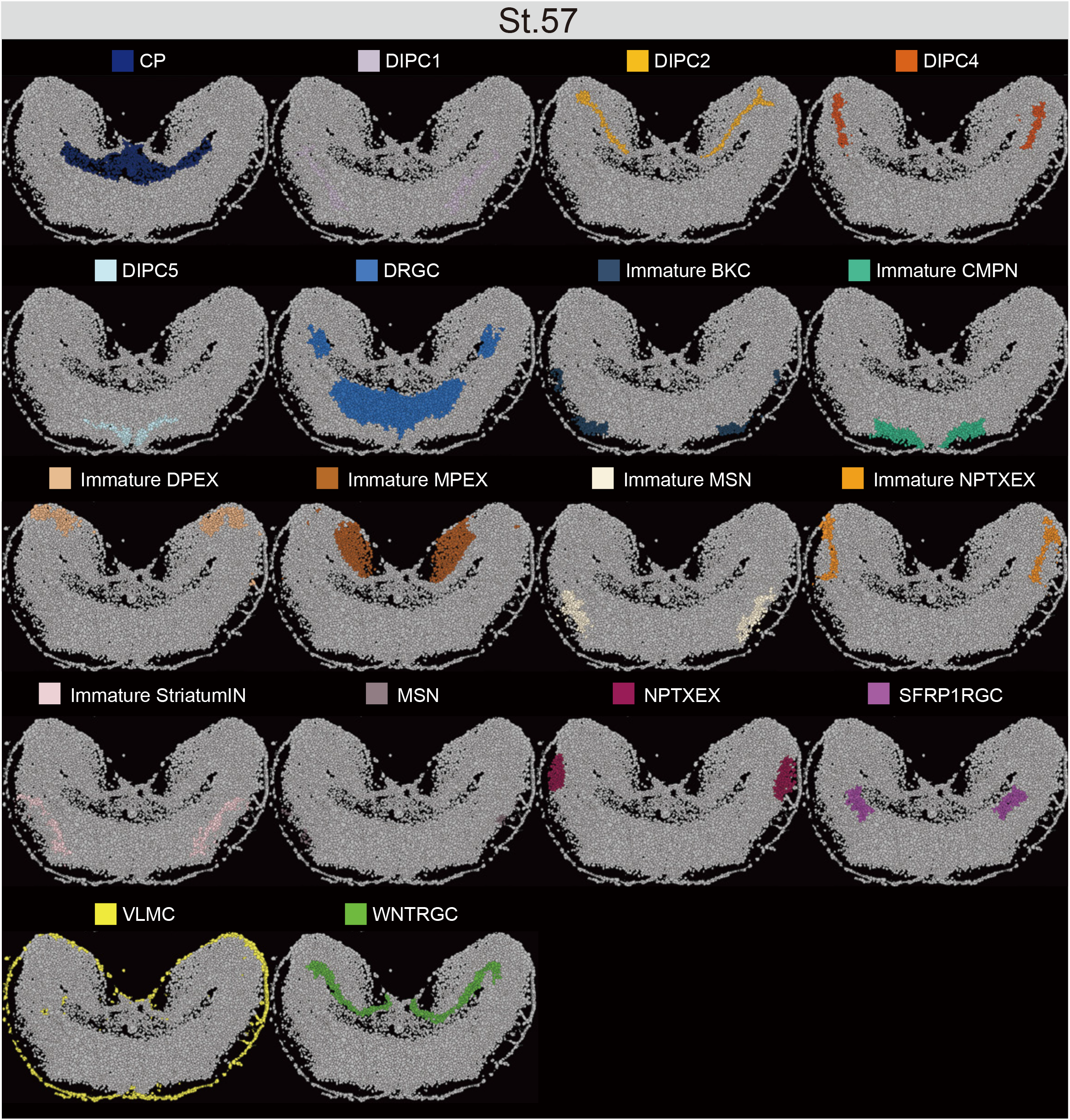
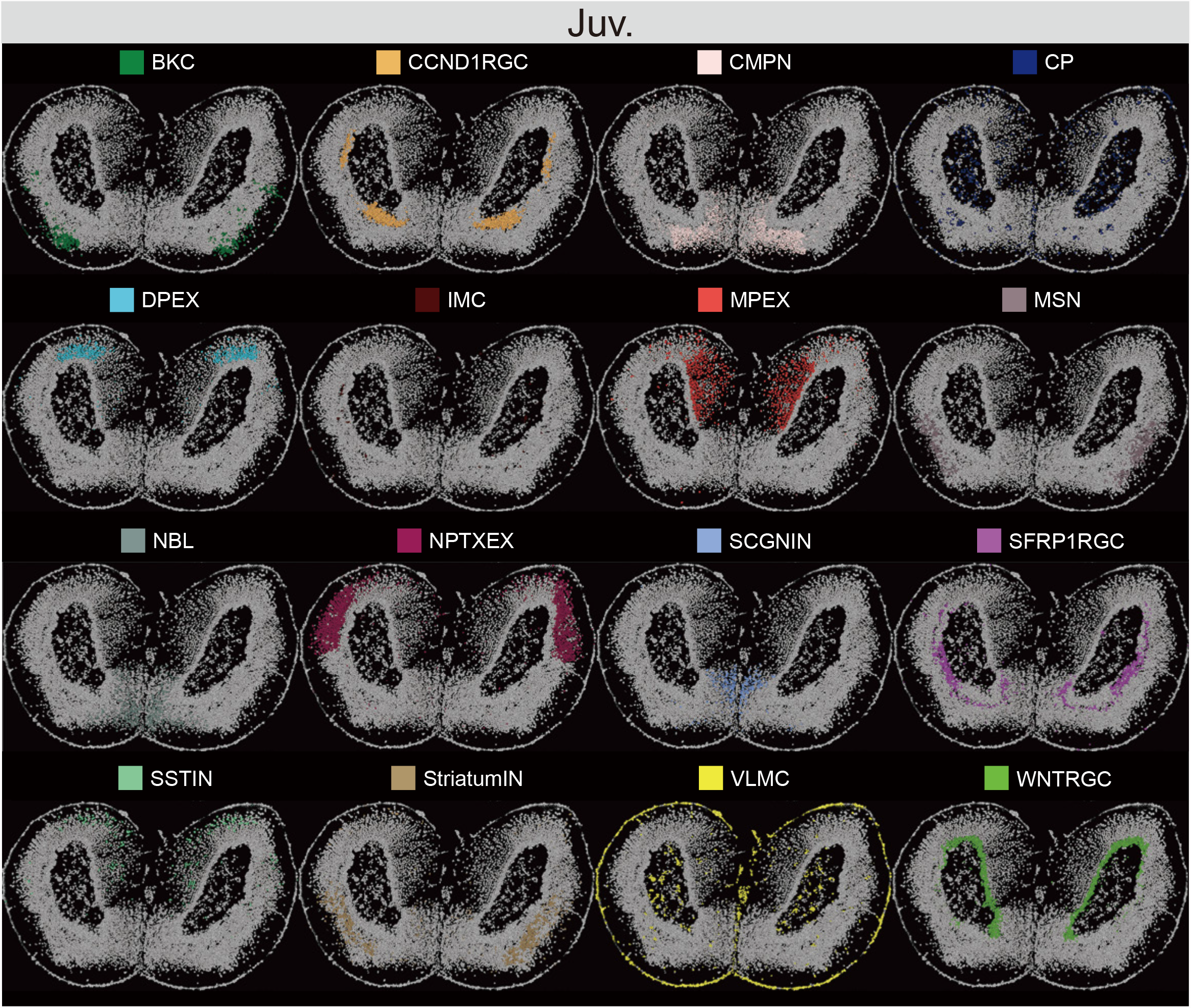
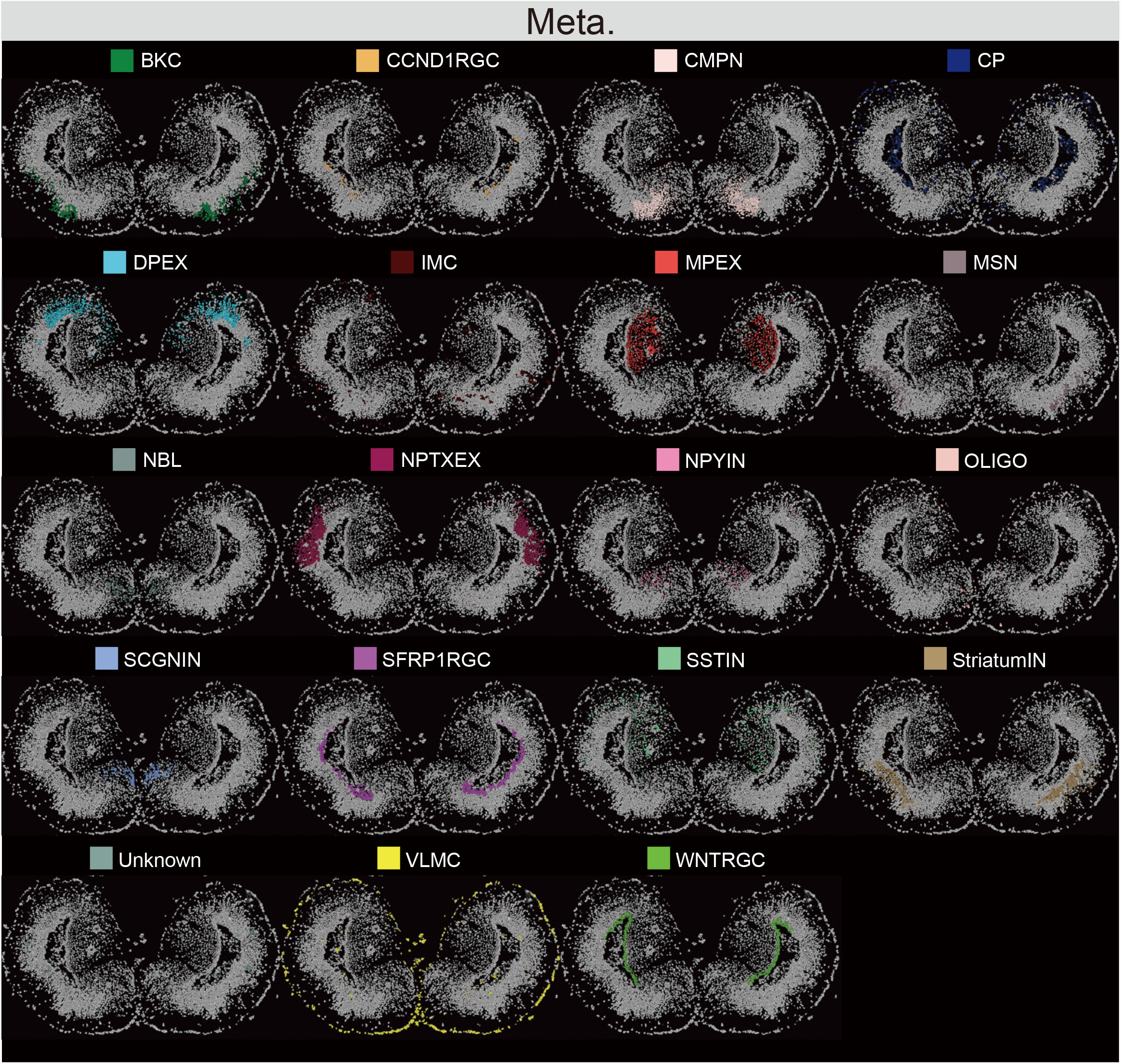
Global spatial profiling of cell types, related to Figure 2. Spatial visualization of cell type distribution in different axolotl telencephalon sections profiled by Stereo-seq in Figure 2A. Cells were colored by their annotation.

**Figure S12.**
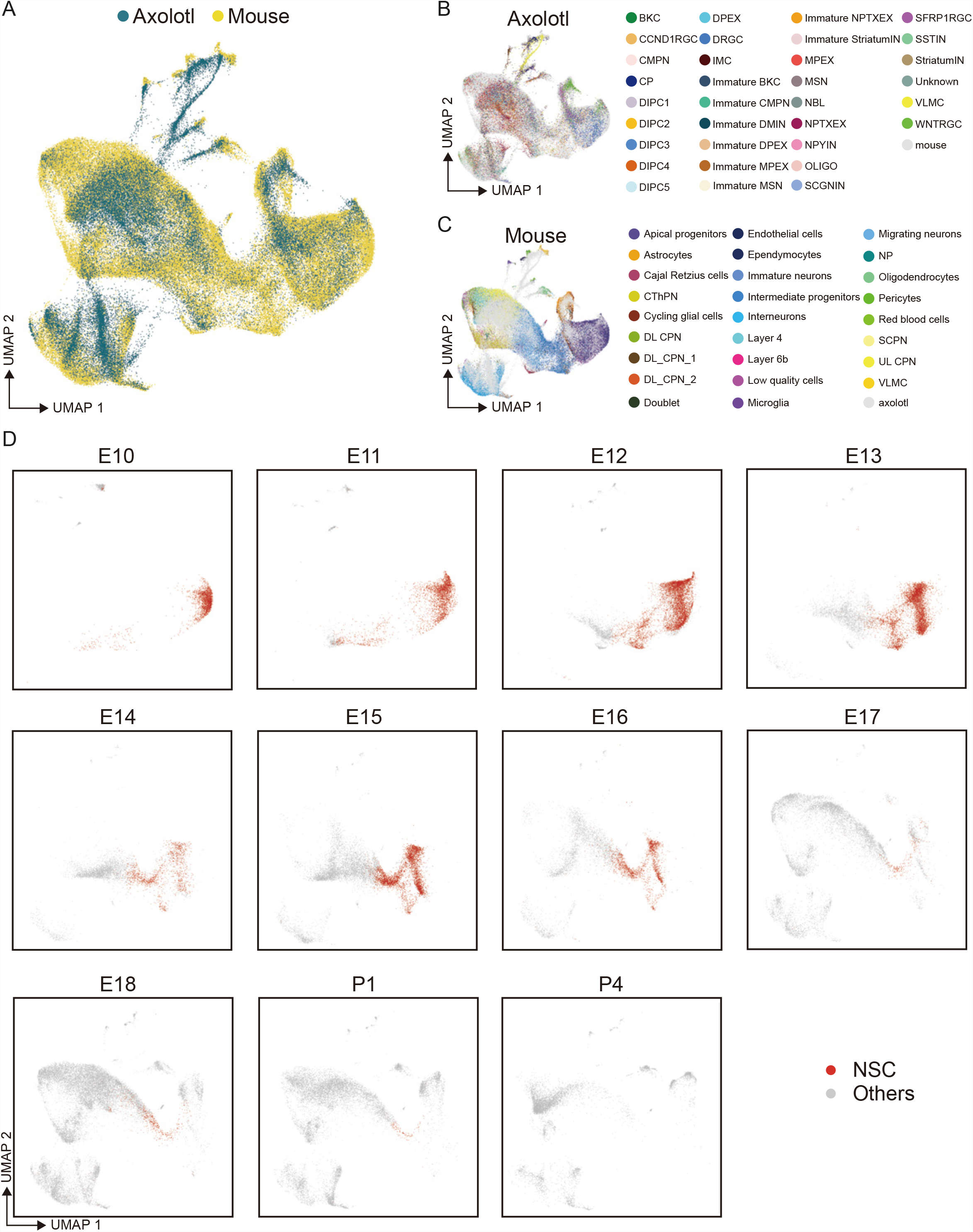
Comparative analysis of NSC between axolotl and mouse, related to Figure 2. (A) UMAP of integrated axolotl Stereo-seq data and mouse single-cell RNA-seq data. (B-C) UMAP visualization of cell types identified in axolotl data (B) and mouse data(C). (D) UMAP visualization of mouse embryonic neural stem cells at each stage profiled by single-cell RNA-seq. Red dots represent neural stem cells at each stage, gray dots represent other cells at respective stage.

**Figure S13.**
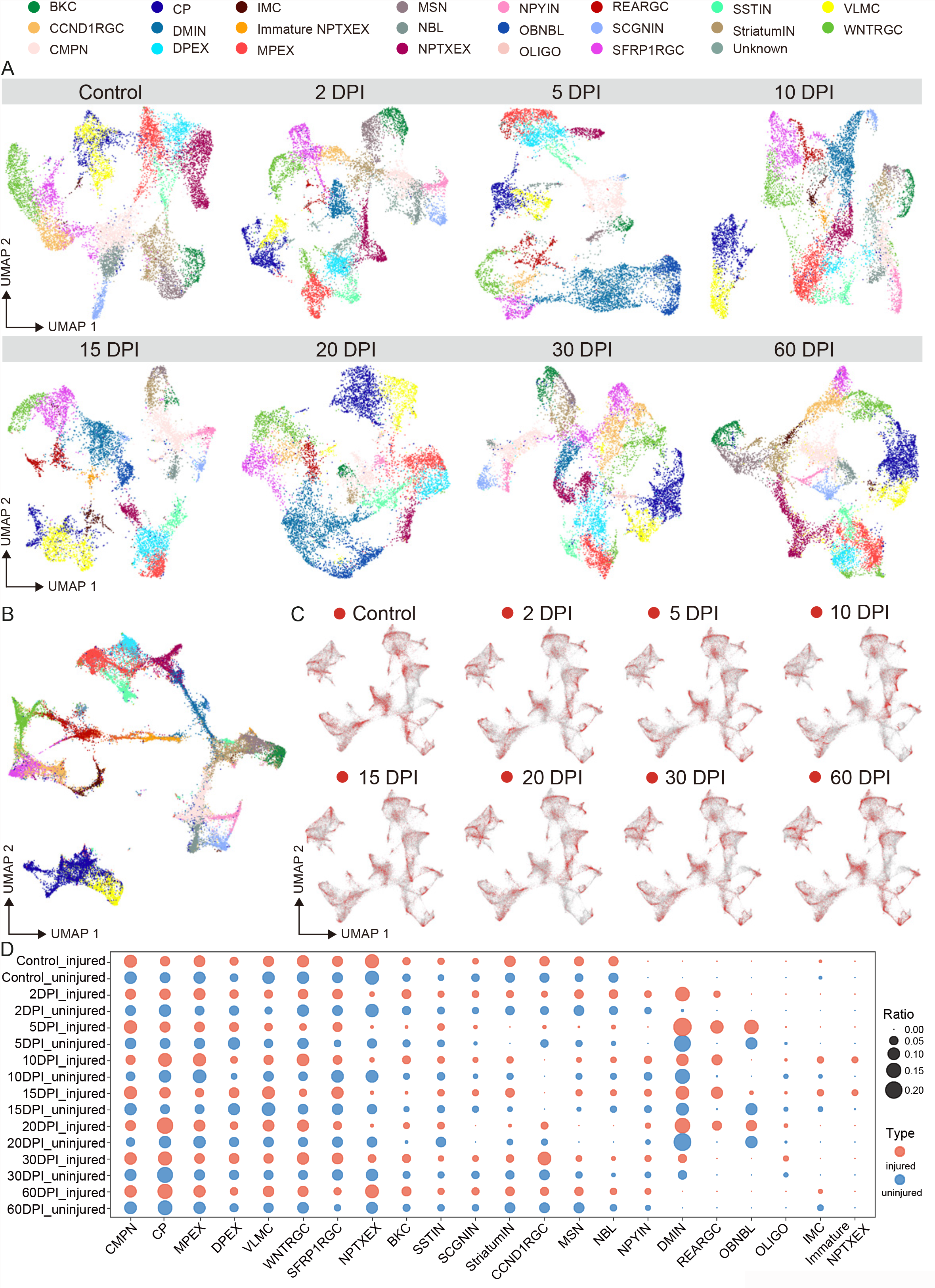
Cell dynamics of axolotl telencephalon regeneration, related to Figure 3. (A) UMAP visualizing the clusters at each stage. 23 cell types are identified and annotated as in Figure 3A. (B) UMAP showing the integration of 23 cell types across different stages. (C) UMAP showing the distribution of cells from each sampling stage in Figure 3A. Red dots represent the cells from the corresponding time point, gray dots represent all the other cells. (D) Dotplot showing the cell ratio dynamics from control to 60 DPI during axolotl telencephalon regeneration. Red dots represent cellular ratio in injured hemisphere, blue dots represent cellular ratio in uninjured hemisphere, dot size reflect the ratio.

**Figure S14.**
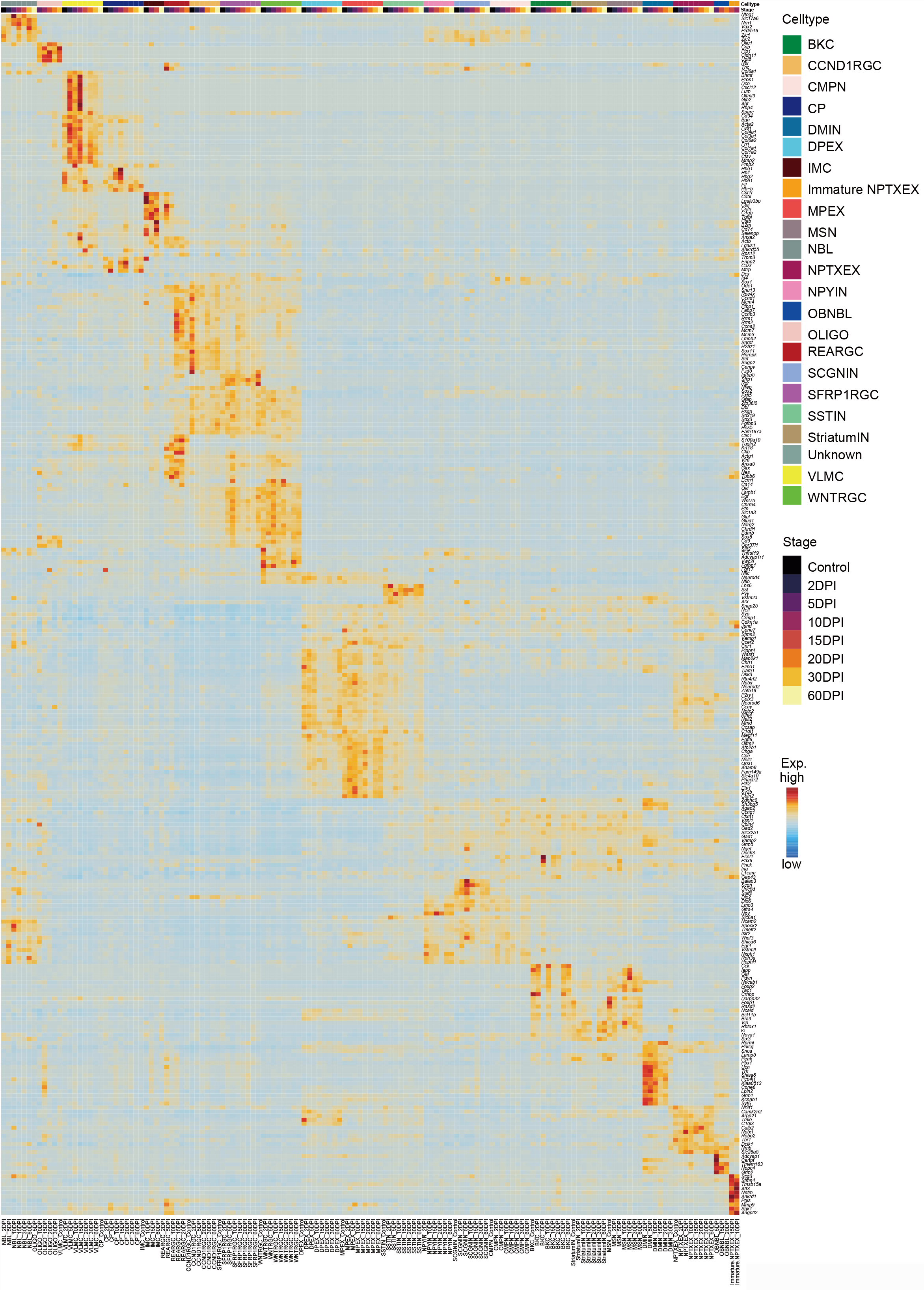
Markers of cell clusters across axolotl telencephalon sections at regenerative stages, related to Figure3. Heatmap showing the normalized expression of marker genes of 23 cell types from axolotl telencephalon sections at regenerative stages profiled by Stereo-seq in Figure 3A.

**Figure S15, S16, S17, S18, S19, S20, S21, S22.**
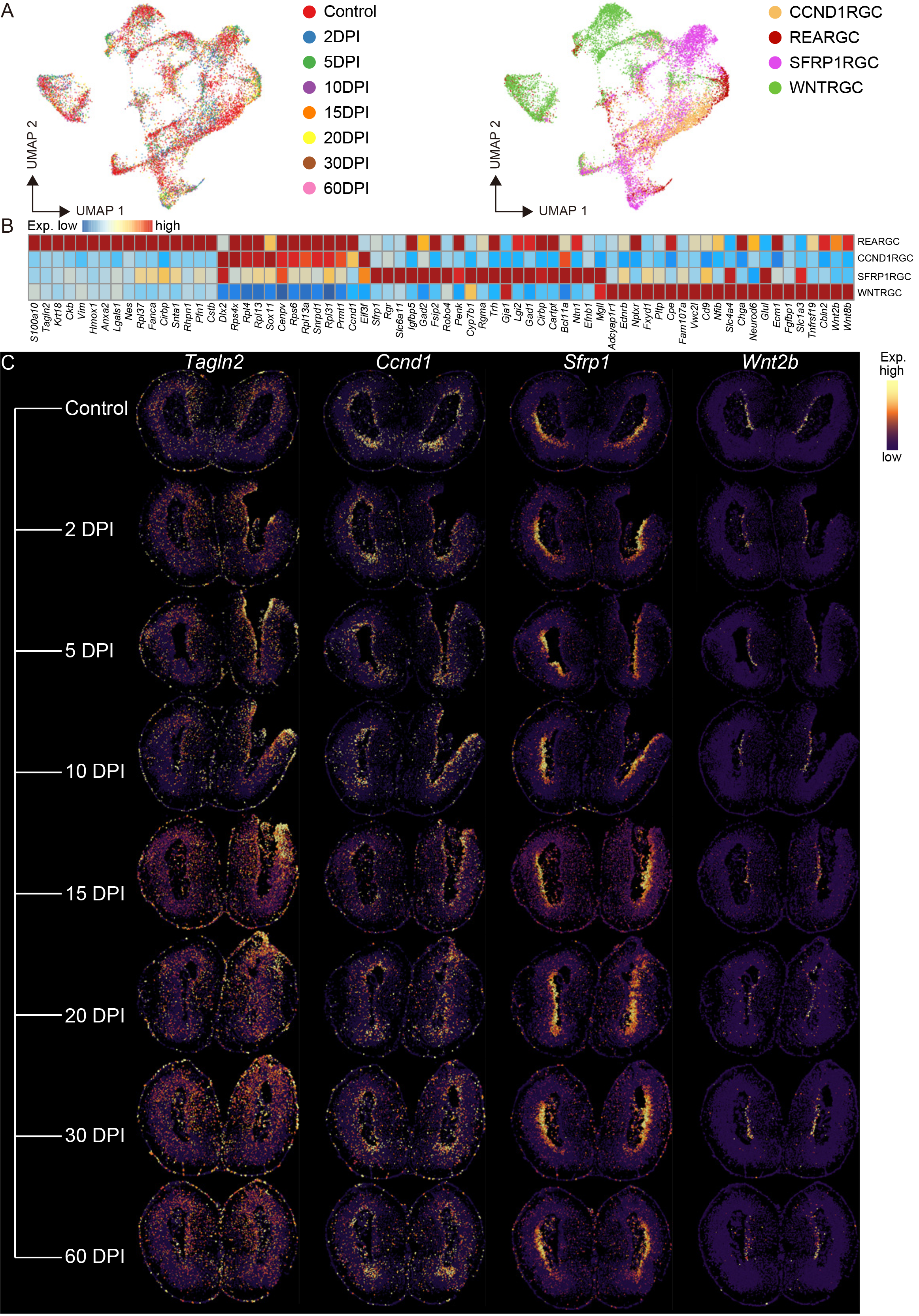
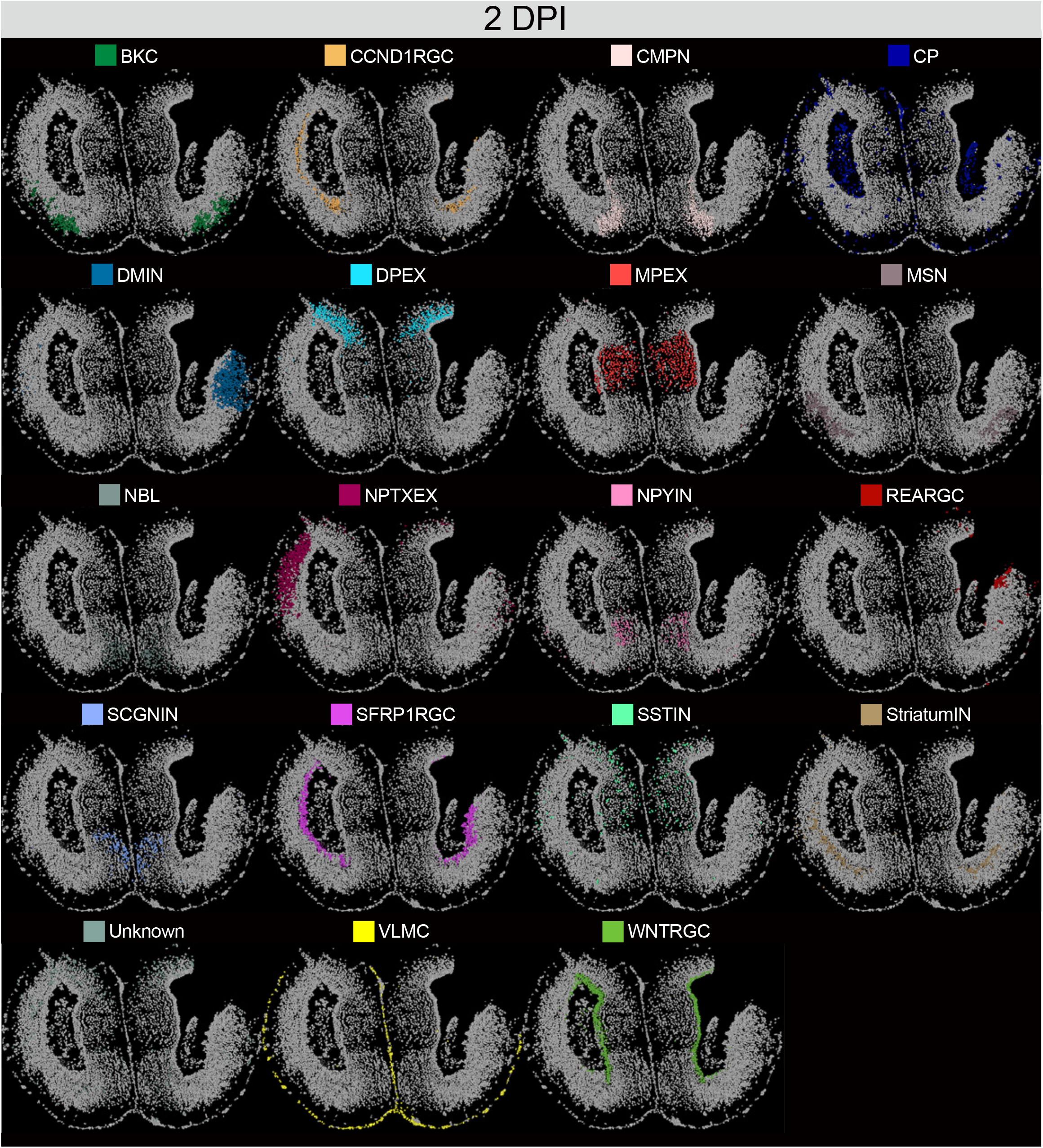
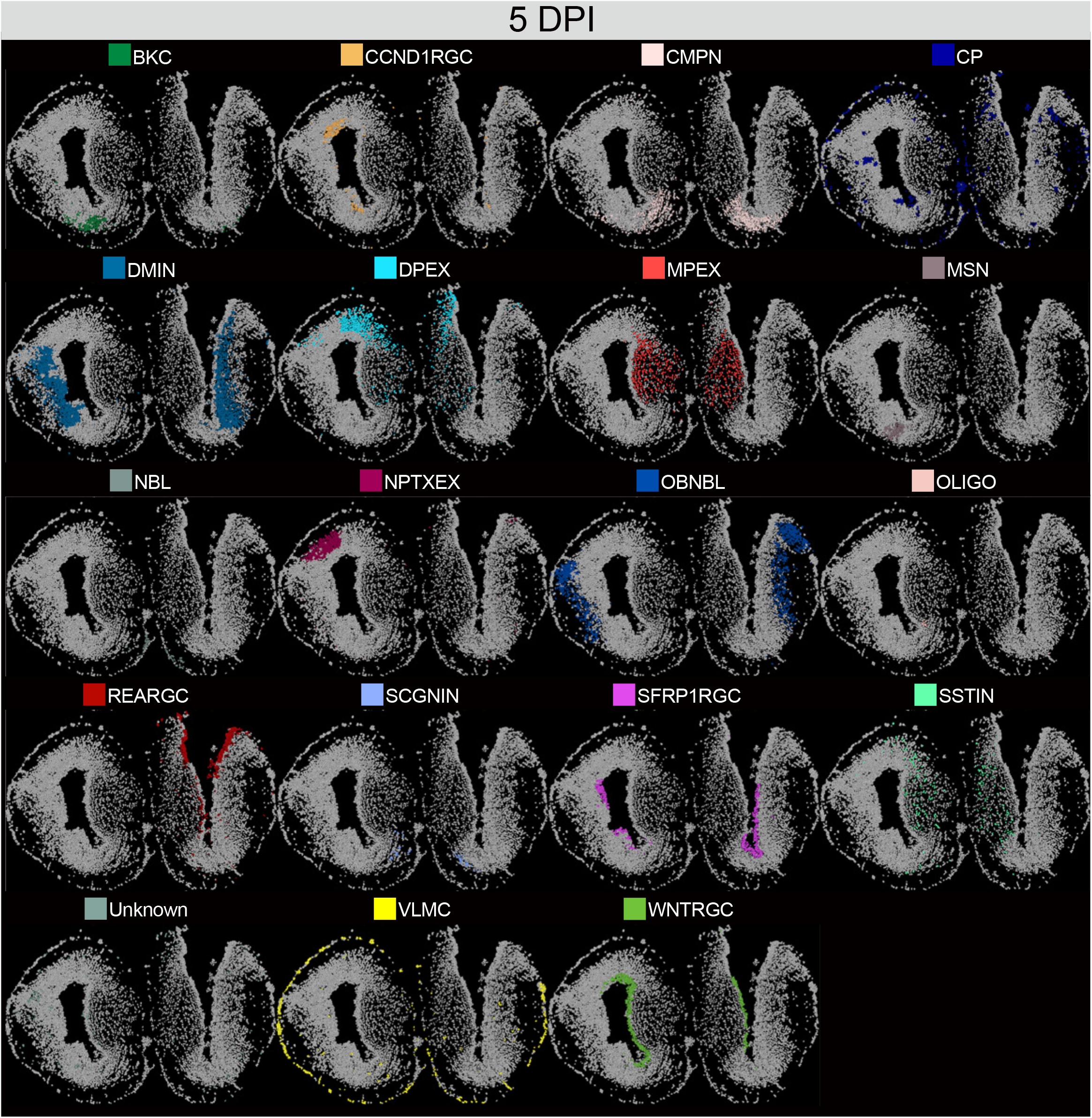
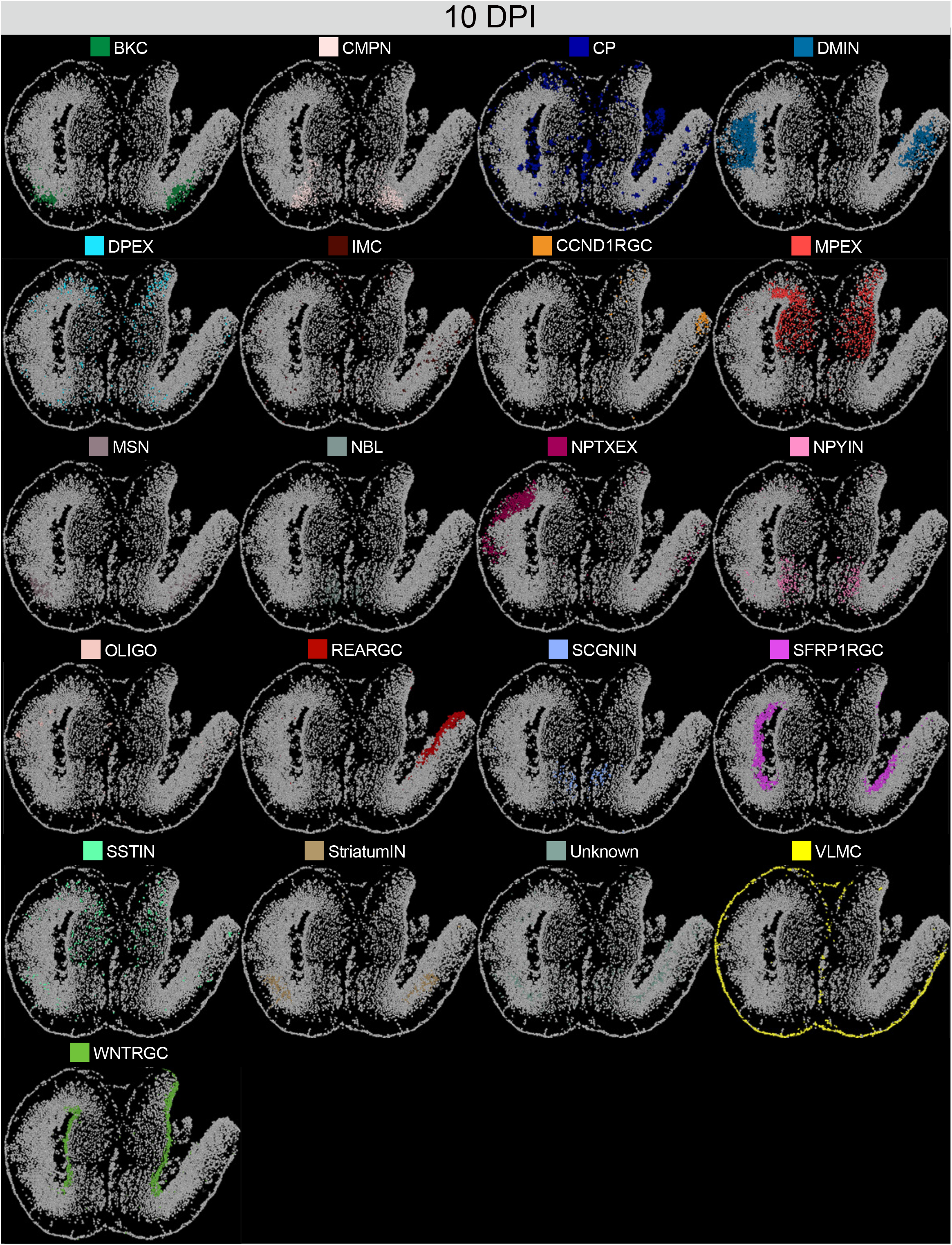
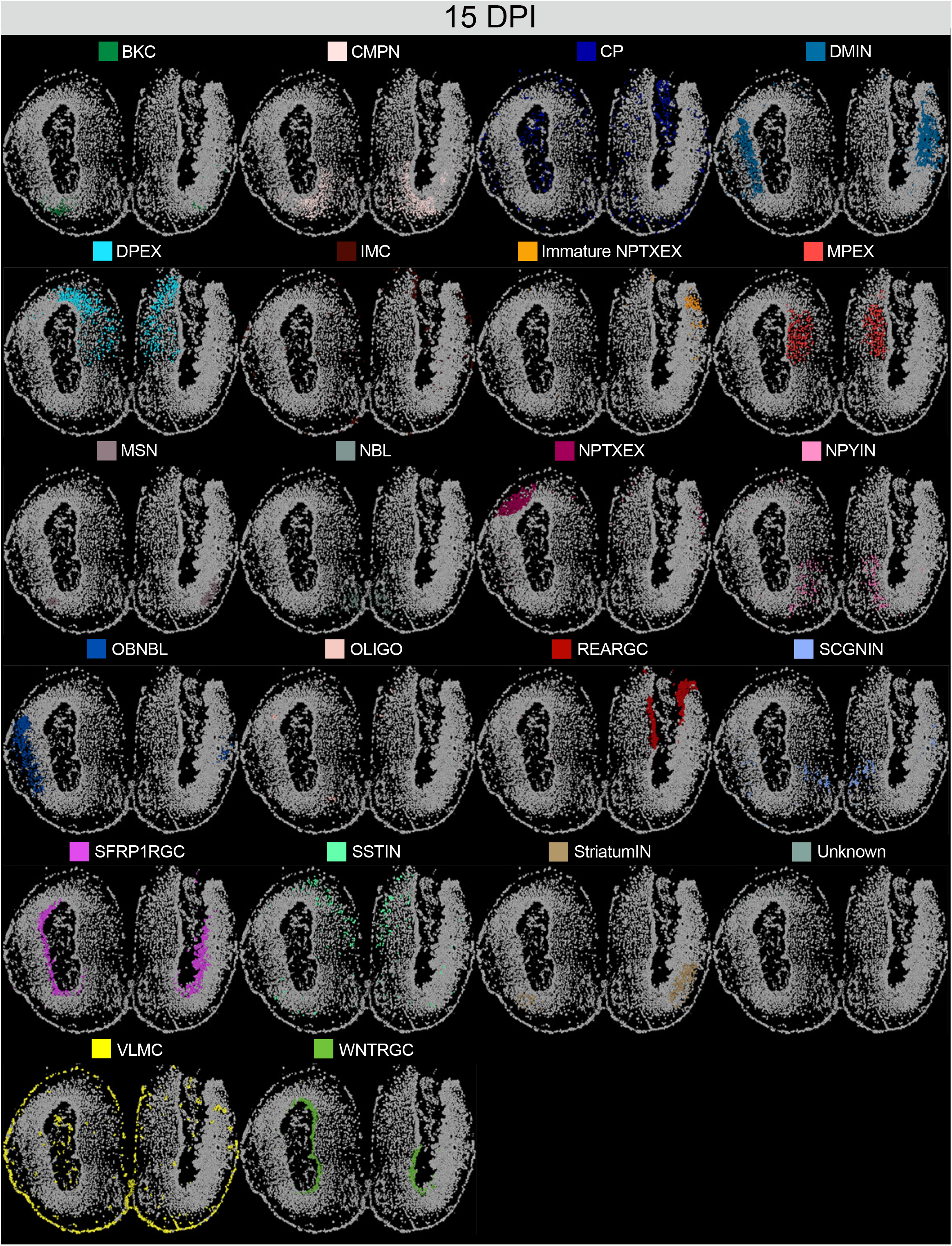
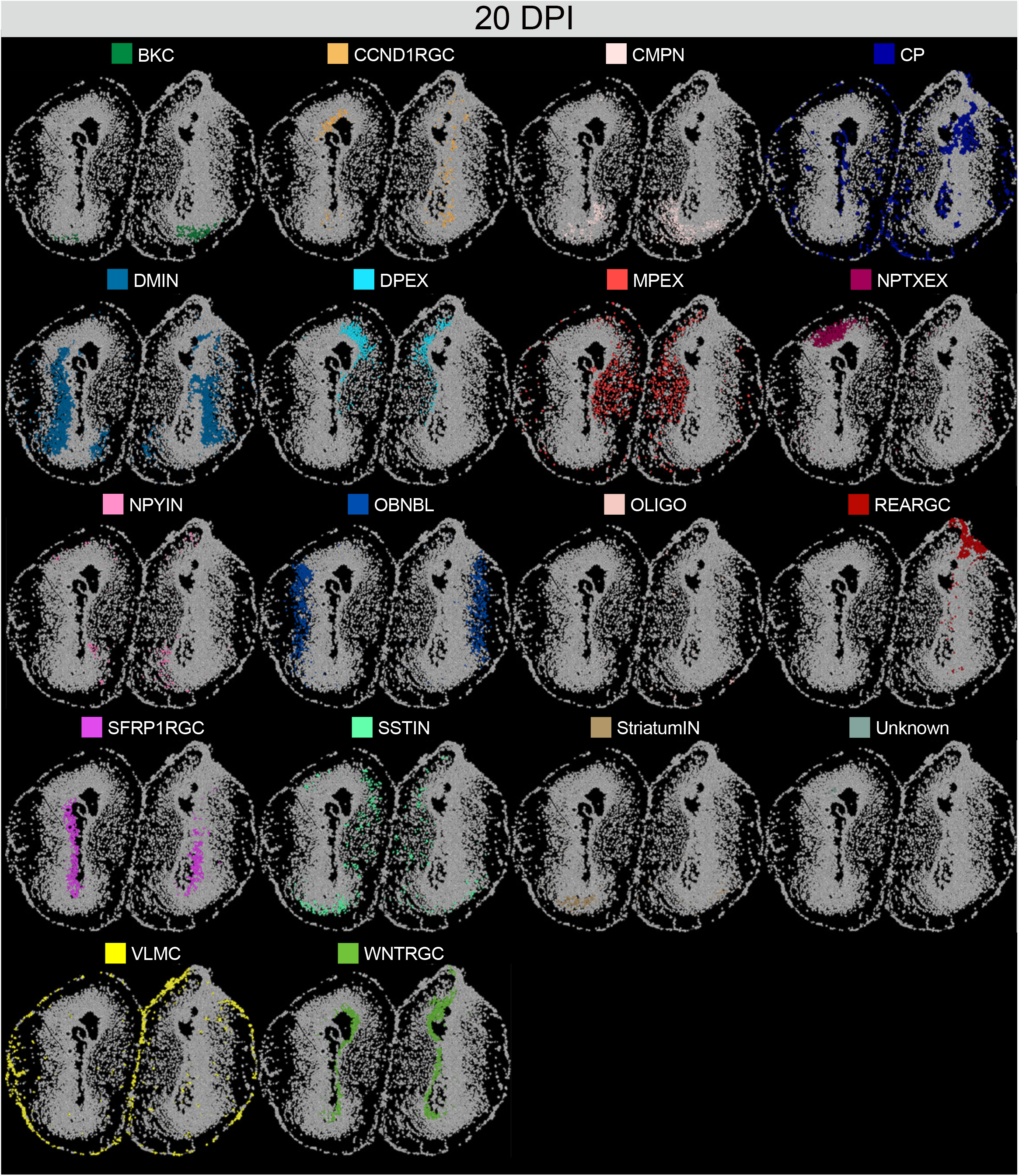
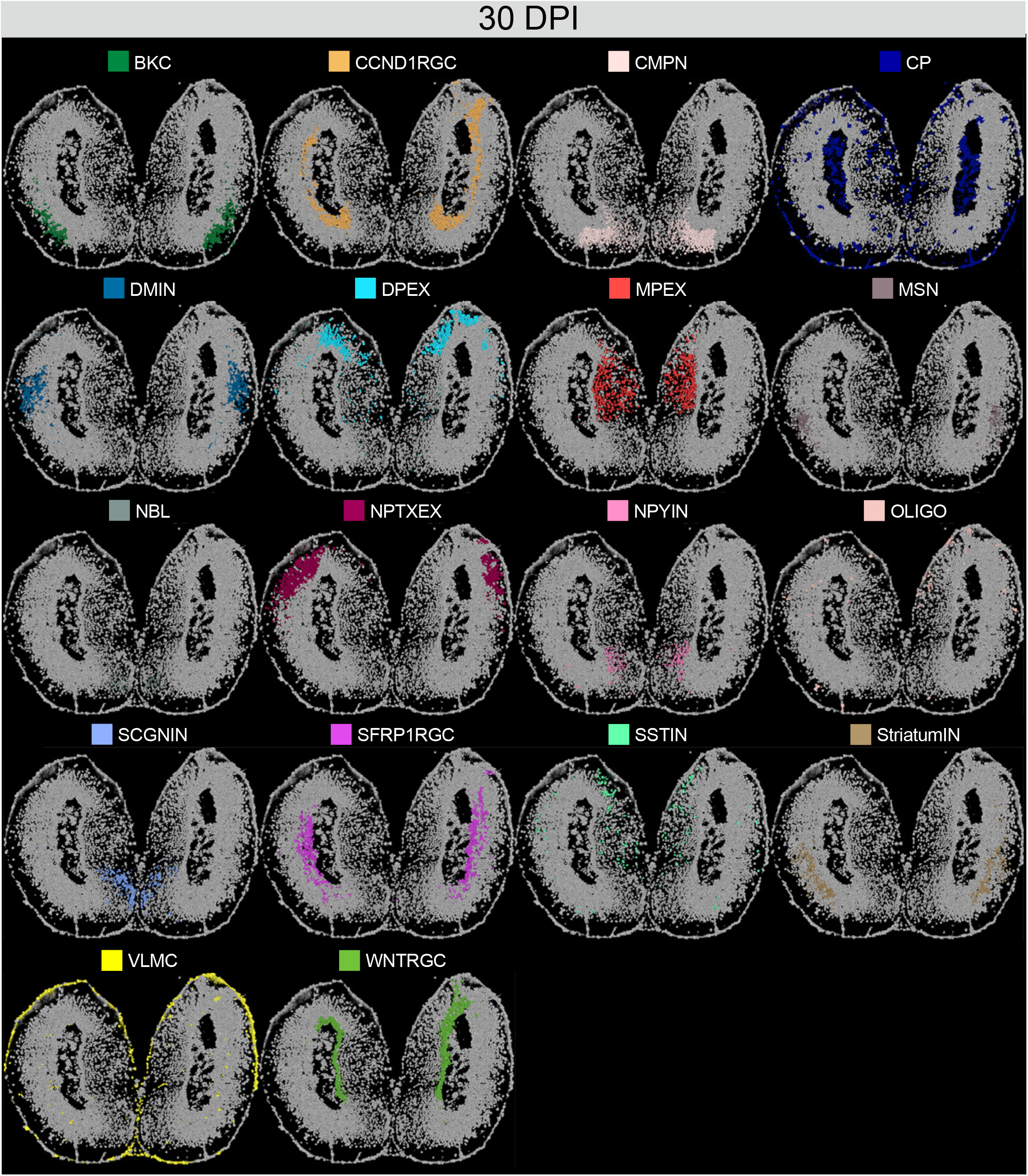
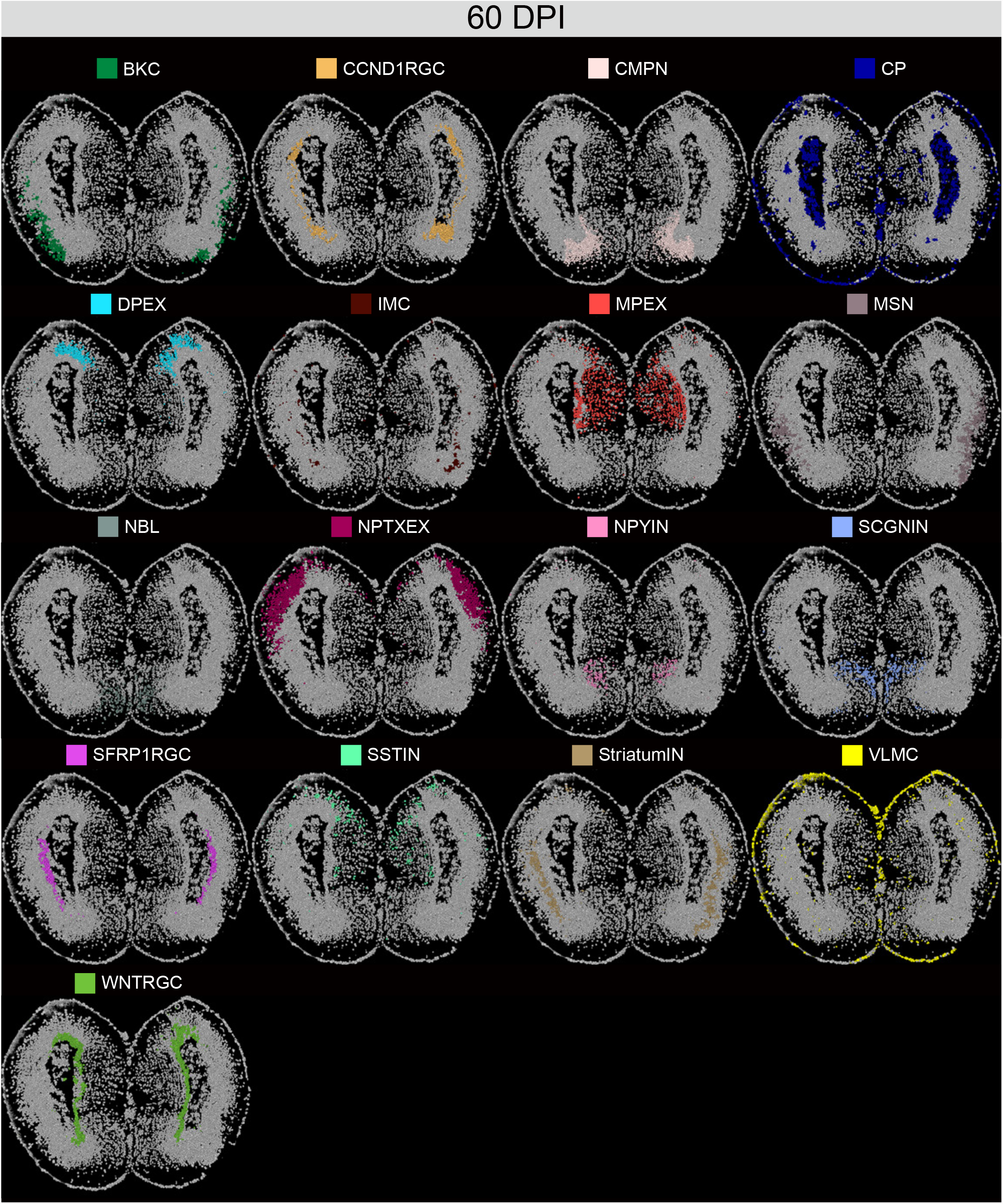
Global spatial profiling of cell types at regenerative stages, related to Figure 3. Spatial visualization of cell type distribution at different axolotl telencephalon regenerative stages profiled by Stereo-seq in Figure 3A. Cells were colored by their annotation.

**Figure S23.**
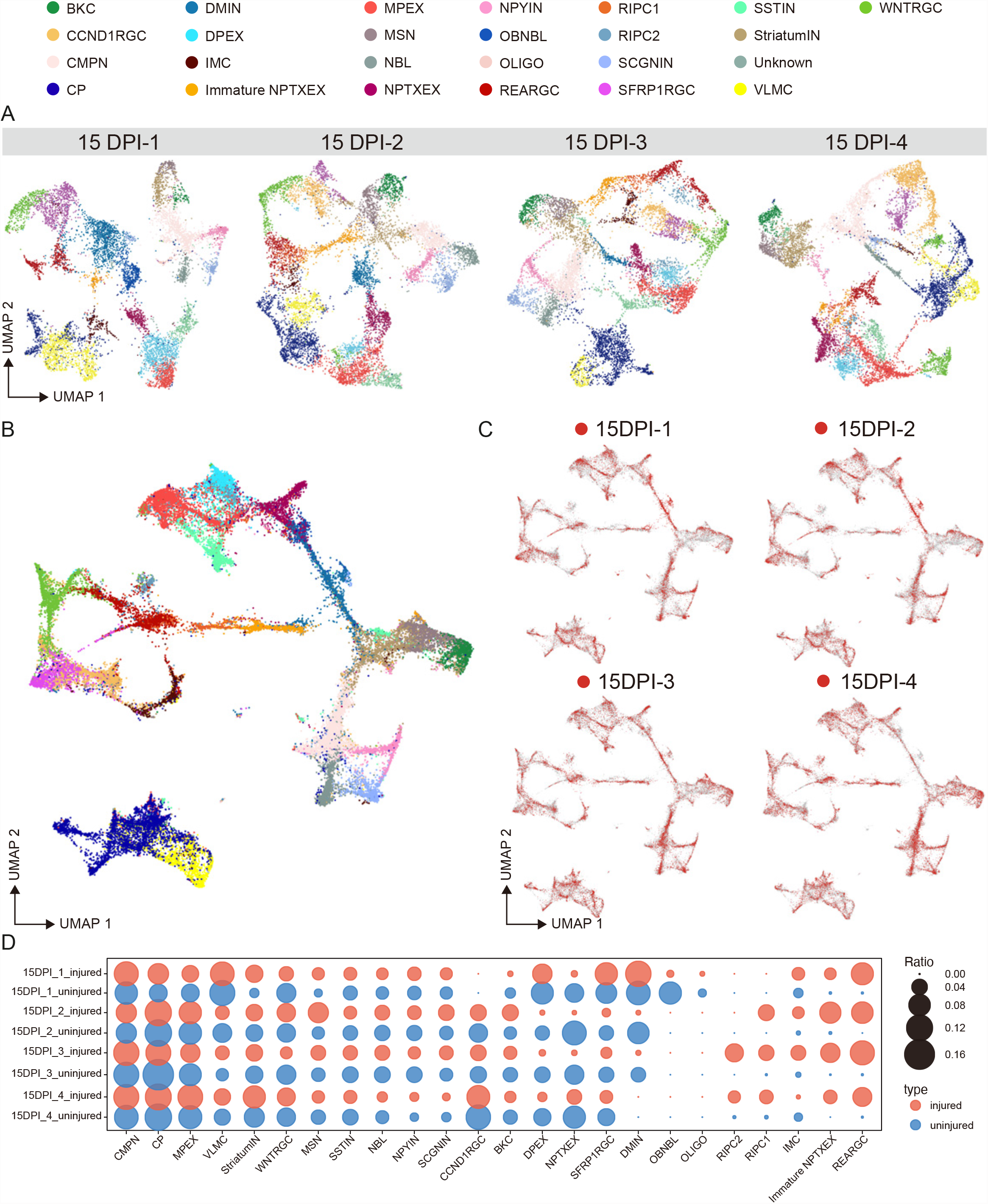
Unsupervised clustering of cells across continuous sections at 15 DPI, related to Figure 4. (A) UMAP visualization of clusters in each section at 15 DPI. 25 cell types are identified and annotated as in Figure 4. (B) UMAP showing the integration of 25 cell types across different sections. (C) UMAP showing the distribution of cells from each sampling section in Figure 4A. Red dots represent the cells from the corresponding time point, gray dots represent all the other cells. (D) Dotplot showing the cell ratio dynamics in different regeneration sections at 15 DPI. Red dots represent cellular ratio in injured hemisphere, blue dots represent cellular ratio in uninjured hemisphere, dot size reflect the ratio.

**Figure S24.**
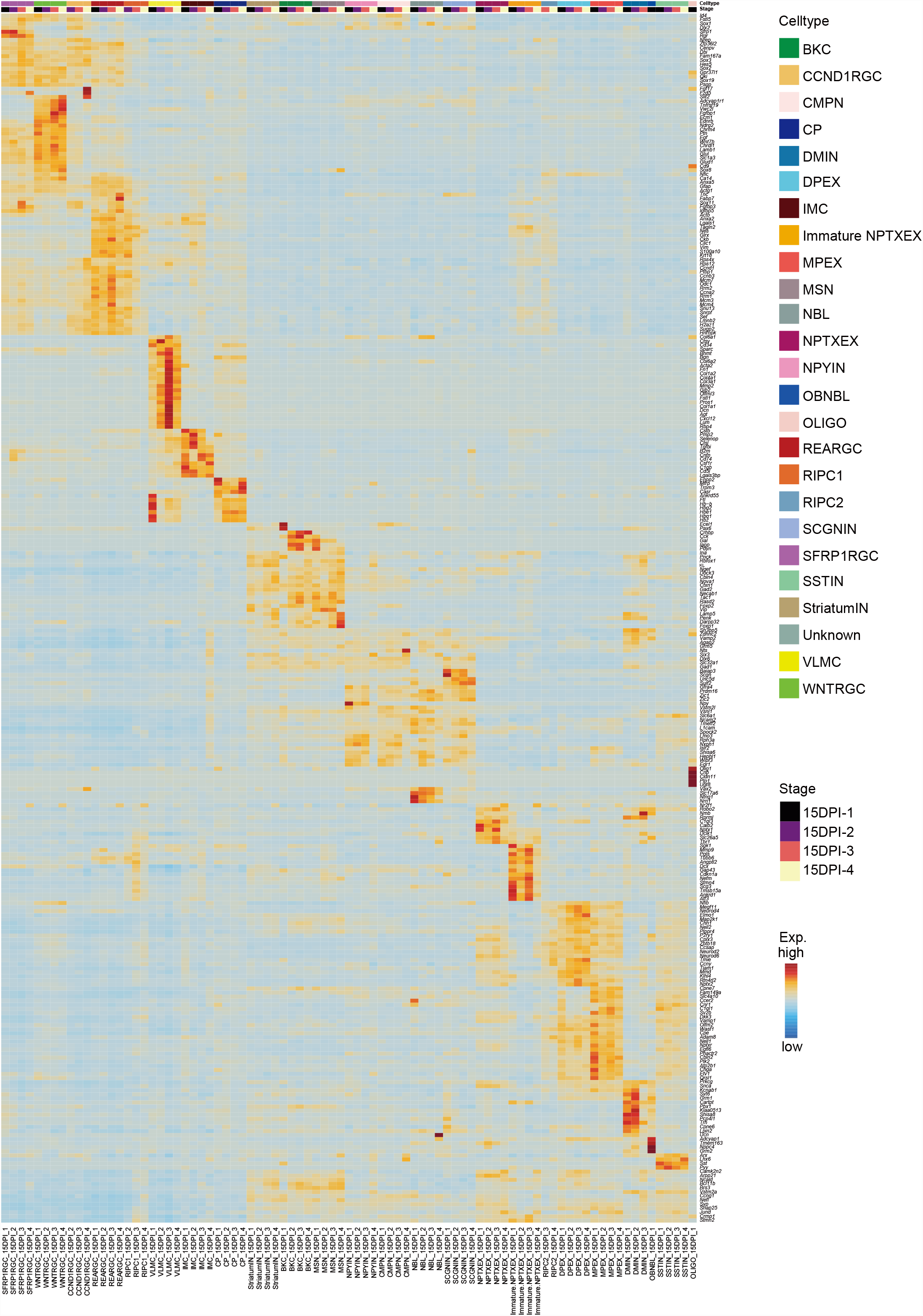
Markers of cell clusters across continuous sections at 15 DPI, related to Figure 4. Heatmap showing the normalized expression of marker genes for 25 cell types in 4 neighbor axolotl telencephalon sections at 15 DPI profiled by Stereo-seq in Figure 4A.

**Figure S25, S26, S27.**
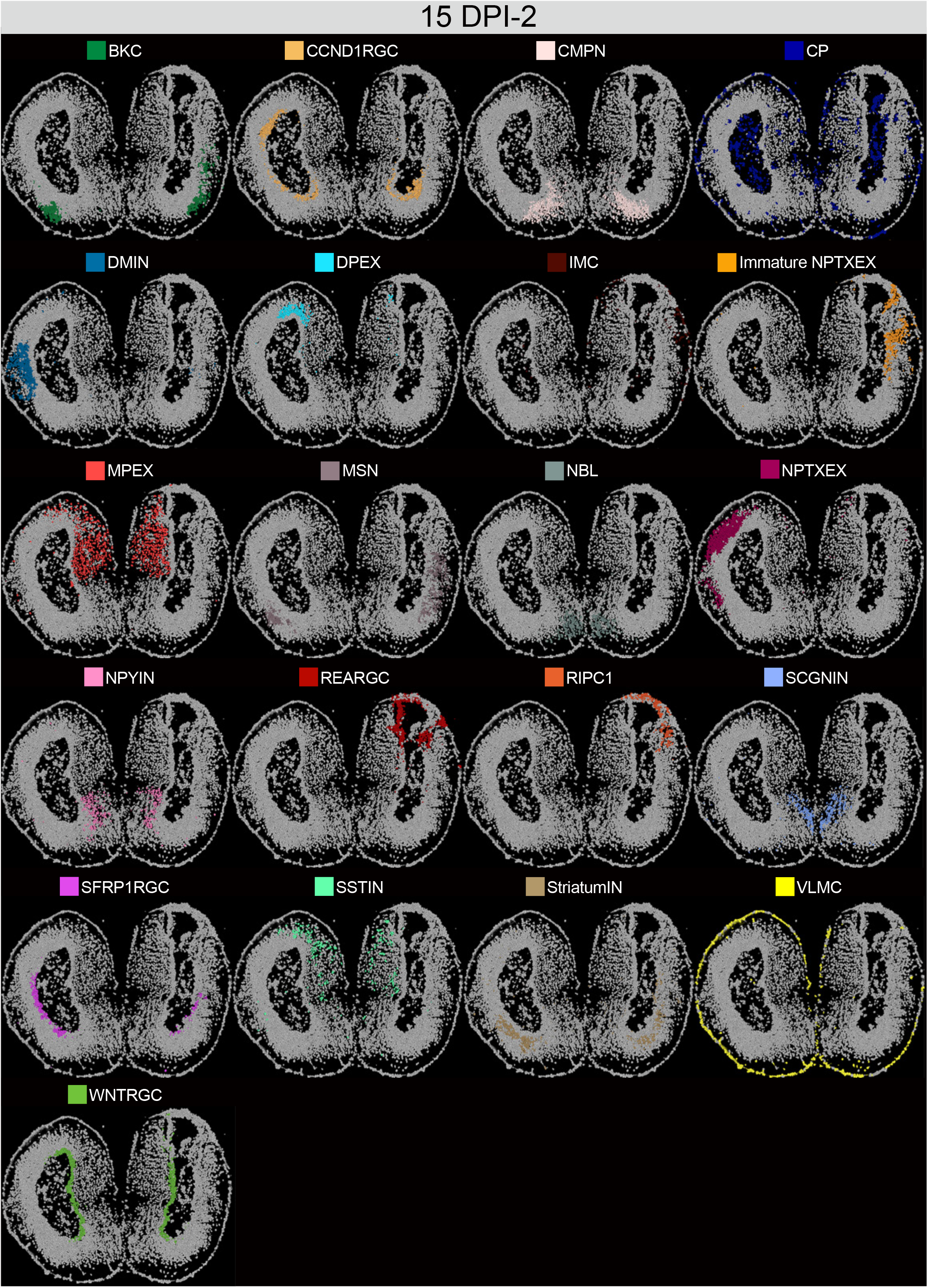
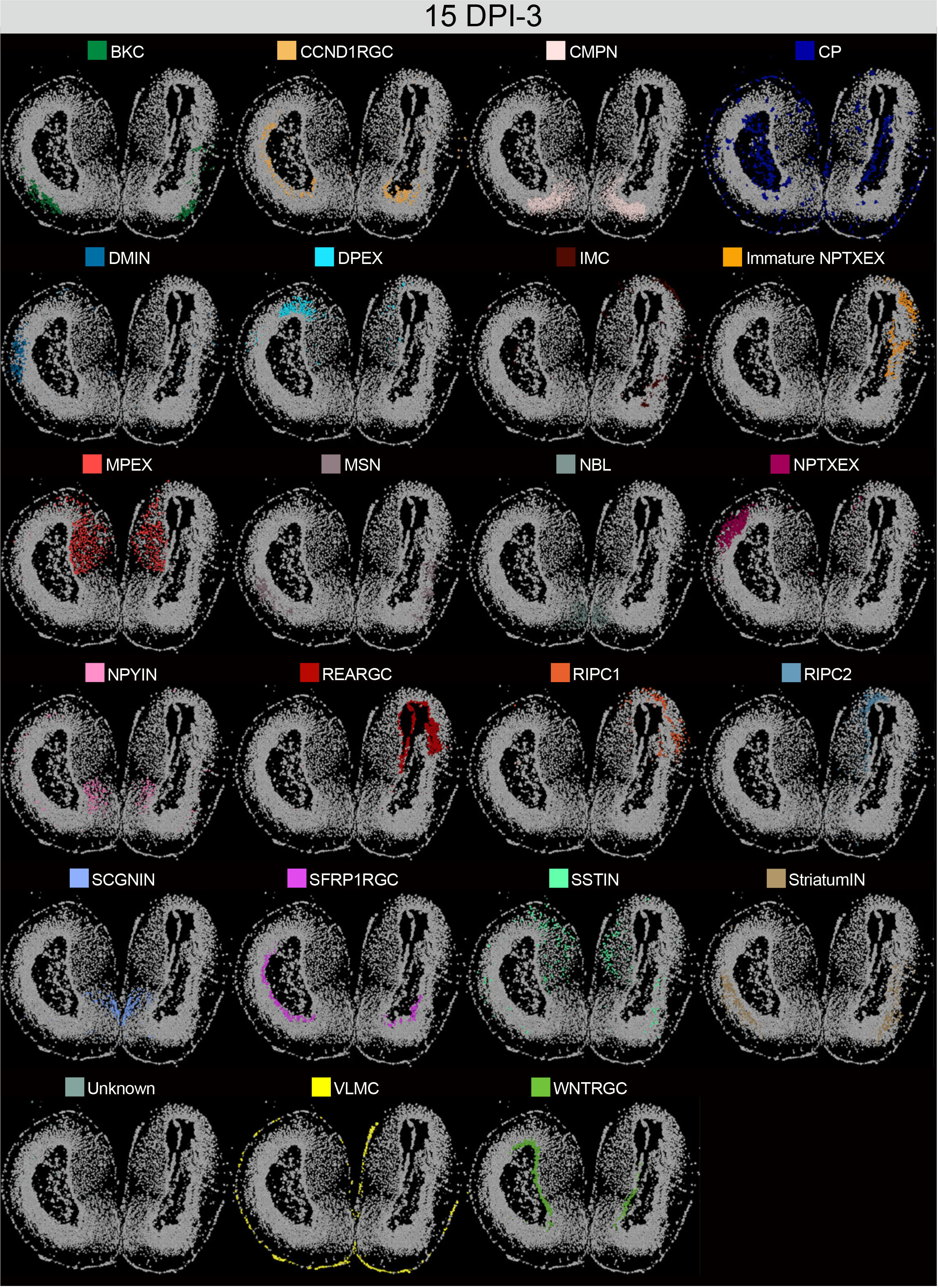
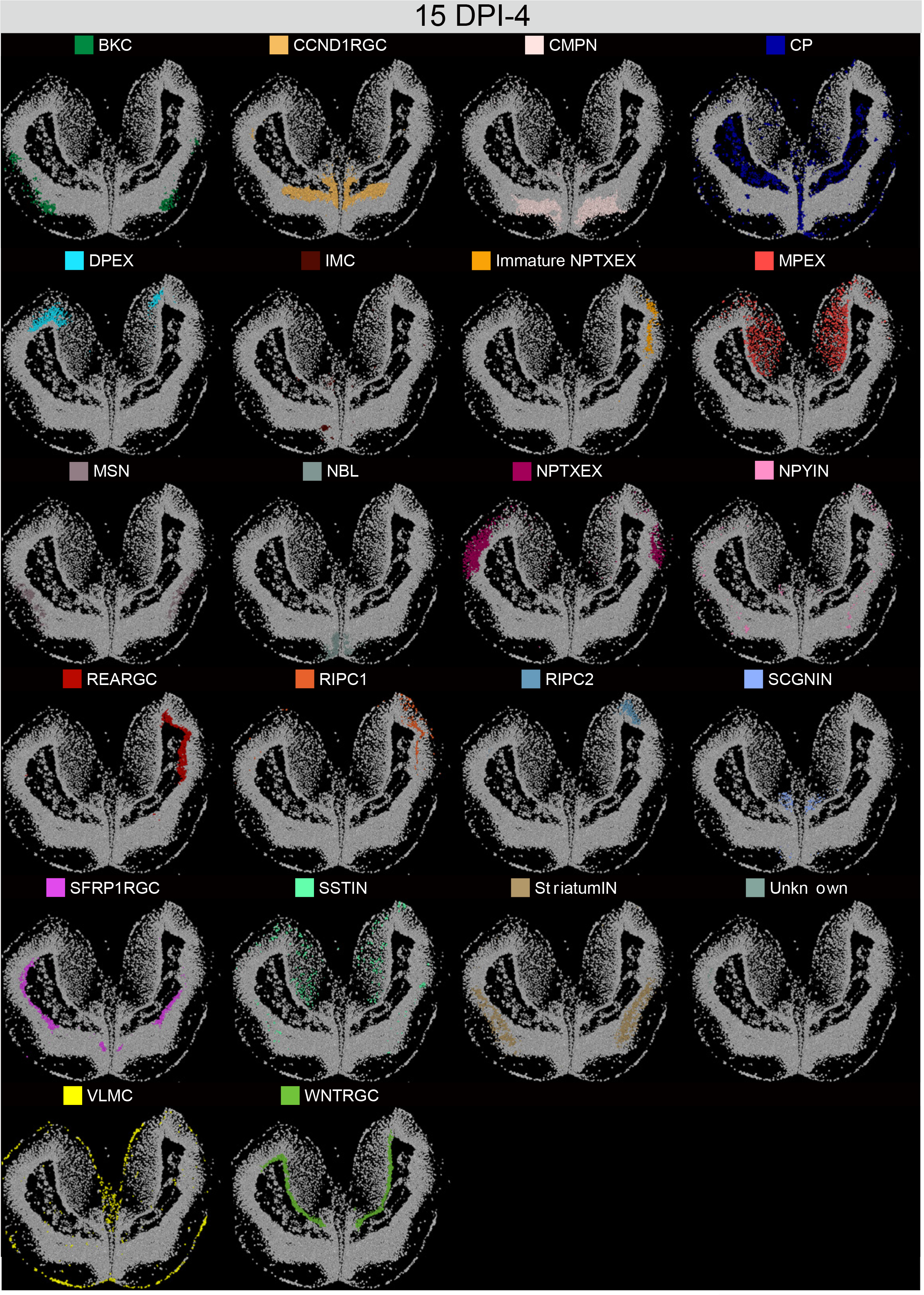
Global spatial profiling of cell types across continuous sections at 15 DPI, related to Figure 4. Spatial visualization of cell type distribution in neighbor regenerating sections of the axolotl telencephalon at 15 DPI profiled by Stereo-seq in Figure 4A. Cells were colored by their annotation.

**Figure S28.**
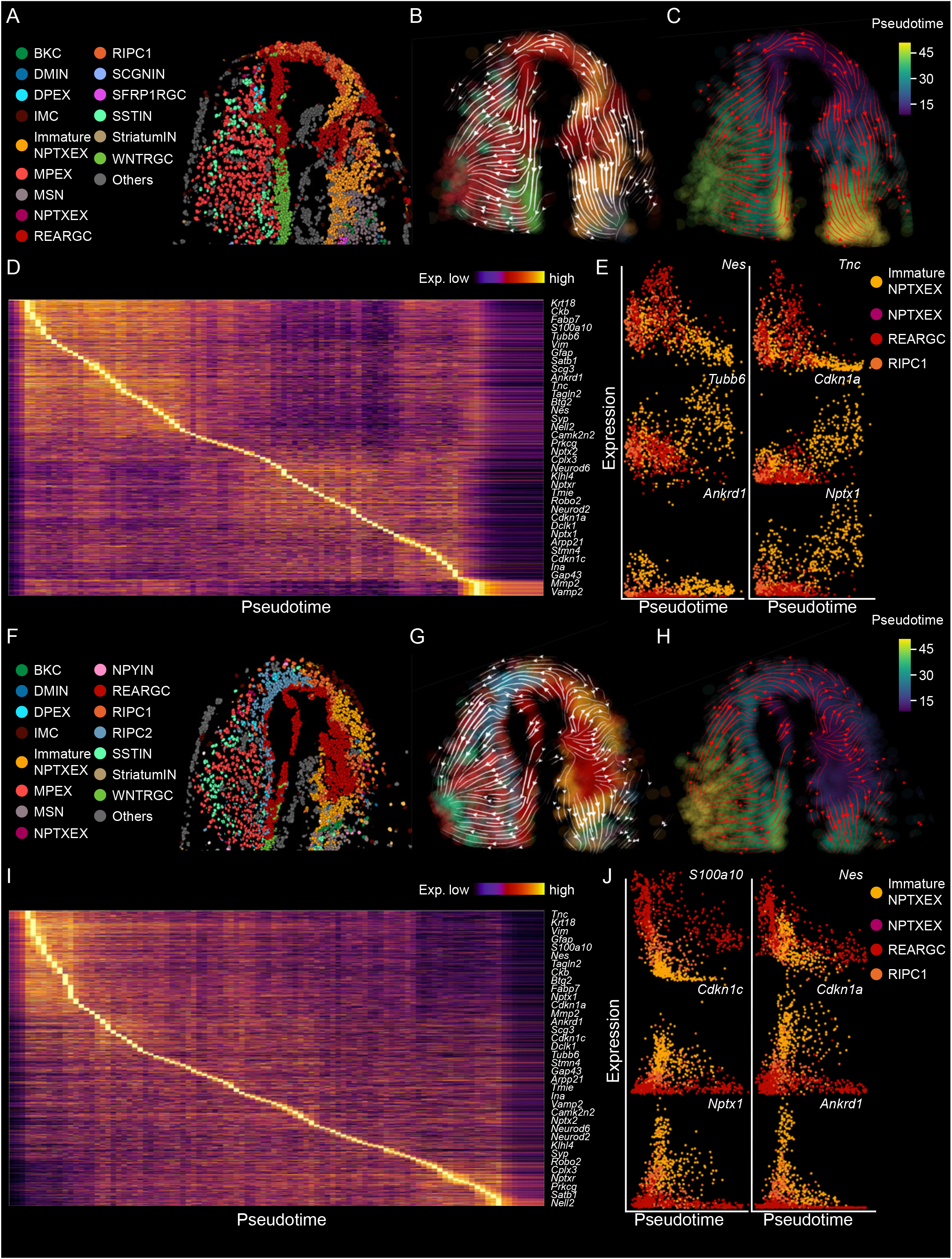
RNA velocity analysis of 15 DPI-2 and 15 DPI-3, related to Figure 4. (A) Spatial visualization of cell types involved in regeneration around the injured site of 15DPI-2(A). (B-C) RNA velocity streamline plots simulating the regeneration trajectory of 15DPI-2. Cells are colored by their annotation(B) or pseudotime (C). (D) Expression heatmap of genes with high transitional activity in a pseudo-temporal order in section 15 DPI-2. (E) Scatter plots showing the pseudotime kinetics of *Nes*, *Tnc*, *Tubb6*, *Cdkn1a*, *Ankrd1* and *Nptx1* in different cell types of 15 DPI-2. (F) Spatial visualization of cell types involved in regeneration around the injured site of 15 DPI-3(F). (G-H) RNA velocity streamline plots simulating the regeneration trajectory of 15 DPI-3. Cells are colored by their annotation (G) or pseudotime (H). (I) Expression heatmap of genes with high transitional activity in a pseudo-temporal order in section15 DPI-3. (J) Scatter plots showing the pseudotime kinetics of *S100a10*, *Nes*, *Cdkn1c*, *Cdkn1a*, *Nptx1*and *Ankrd1* in different cell types of 15 DPI-3.

**Figure S29.**
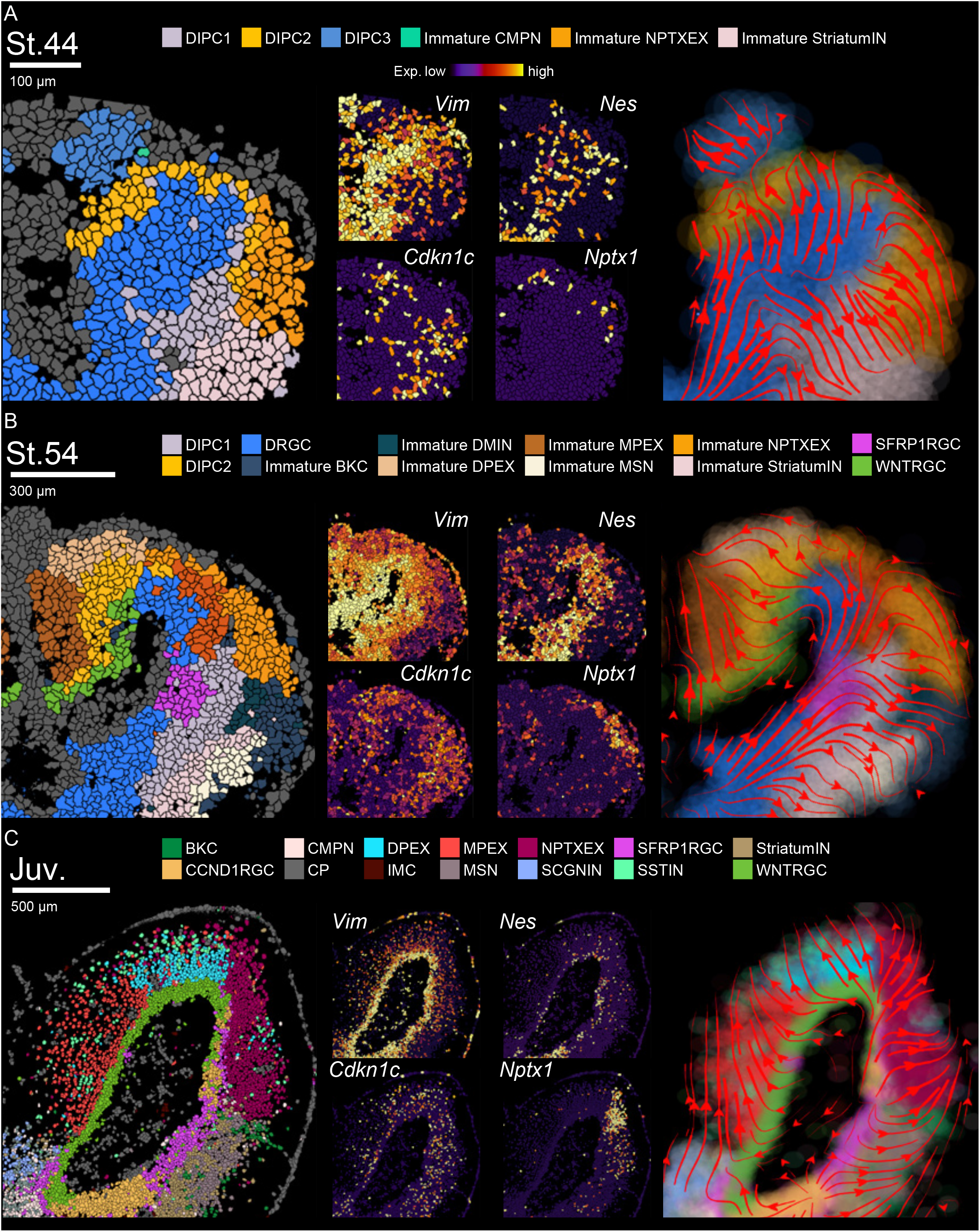
Neurogenesis trajectory analysis of axolotl telencephalon at different developmental stages. (A-C) Spatial visualization of cells involved in telencephalon development at stage 44 (A), stage 54 (B) and juvenile (C) (left). Expression of key marker genes at stage 44 (A), stage 54 (B) and juvenile (C) (middle). RNA velocity streamline plots showing the predicted lineage transition trajectory at stage 44 (A), stage 54 (B) and juvenile (C) (right).

**Figure S30.**
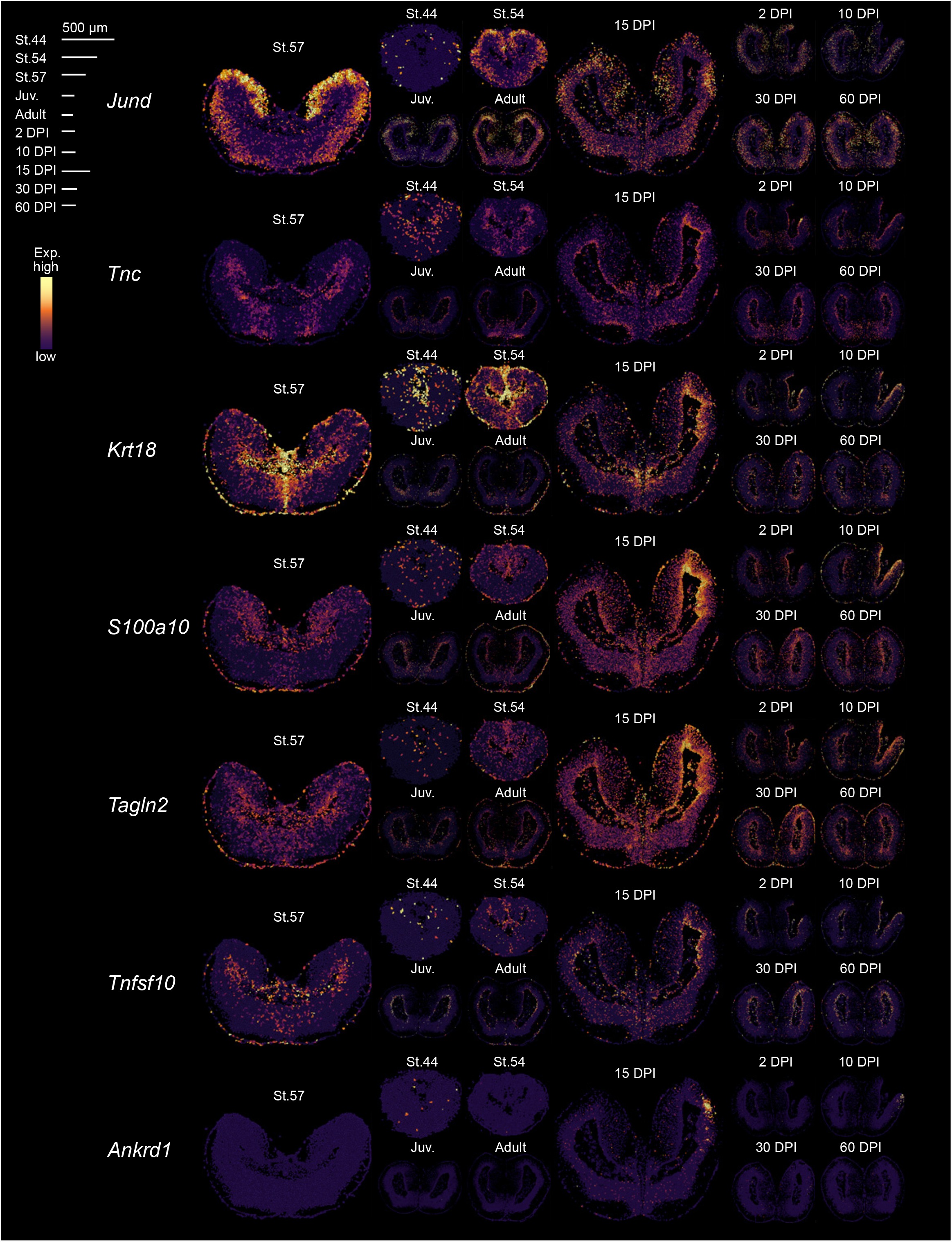
DEGs between development and regeneration. Spatial visualization of the expression of *JunD, Tnc, Krt18*, *S100a10*, *Tagln2, Tnfsf10* and *Ankrd1* in sections of stage 57 and 15 DPI-4.

## TABLE LEGENDS

**Table S1. Description of all samples profiled, cell types and top 20 marker genes.**

**Table S2. Gene list for cluster annotation.**

**Table S3. Gene list of three gene modules defining NSC, cell cycle and translation.**

**Table S4. Gene list of two gene modules in REARGC from 2 to 20 DPI.**

**Table S5. Gene list of the eight patterned groups during development and regeneration processes.**

**Table S6. DEGs of left dorsal pallium region between regeneration and development.**

**Table S7. Primer sequences for ISH.**

